# Functional characterization of the Co^2+^ transporter AitP in *Sinorhizobium meliloti*: a new player in Fe^2+^ homeostasis

**DOI:** 10.1101/2022.09.08.507232

**Authors:** Paula Mihelj, Isidro Abreu, Tomás Moreyra, Manuel González-Guerrero, Daniel Raimunda

**Author notes:** Department of Biology, University of Oxford, Oxford, UK.

## Abstract

Co^2+^ induces the increase of the labile-Fe pool (LIP) by Fe-S cluster damage, heme synthesis inhibition and “free” iron import, which affects cell viability. The N_2_-fixing bacteria, *Sinorhizobium meliloti*, is a suitable model to determine the roles of Co^2+^-transporting Cation diffusion facilitator exporters (Co-eCDF) in Fe^2+^ homeostasis because it has a putative member of this sub-family, AitP, and two specific Fe^2+^-export systems. An insertional mutant of AitP showed Co^2+^ sensitivity and accumulation, Fe accumulation and hydrogen peroxide sensitivity, but not Fe^2+^ sensitivity, despite AitP being a *bona fide* low affinity Fe^2+^ exporter as demonstrated by the kinetic analyses of Fe^2+^ uptake into everted membrane vesicles. Suggesting concomitant Fe^2+^-dependent induced stress, Co^2+^ sensitivity was increased in strains carrying mutations in AitP and Fe^2+^ exporters which did not correlate with the Co^2+^ accumulation. Growth in the presence of sub-lethal Fe^2+^ and Co^2+^ concentrations suggested that free Fe-import might contribute to Co^2+^ toxicity. Supporting this, Co^2+^ induced transcription of Fe-import system and genes associated with Fe homeostasis. Analyses of total protoporphyrin content indicates Fe-S cluster attack as the major source for LIP. AitP-mediated Fe^2+^-export is likely counterbalanced via a non-futile Fe^2+^-import pathway. Two lines of evidence support this: i) an increased hemin uptake in presence of Co^2+^ was observed in WT *vs*. AitP mutant, and ii) hemin reversed the Co^2+^ sensitivity in the AitP mutant. Thus, the simultaneous detoxification mediated by AitP aids cells to orchestrate an Fe-S cluster salvage response, avoiding the increase in the LIP caused by the disassembly of Fe-S clusters or free iron uptake.

**Importance:** Cross-talk between iron and cobalt has been long recognized in biological systems. This is due to the capacity of cobalt to interfere with proper iron utilization. Cells can detoxify cobalt by exporting mechanisms involving membrane proteins known as exporters. Highlighting the cross-talk, the capacity of several cobalt exporters to also export iron is emerging. Although biologically less important than Fe^2+^, Co^2+^ induces toxicity by promoting intracellular Fe release, which ultimately causes additional toxic effects. In this work, we describe how the N_2_-fixating rhizobial cells solve this perturbation by clearing Fe through a Co^2+^-exporter, in order to reestablish intracellular Fe-levels by importing non-free Fe, heme. This piggyback-ride type of transport may aid bacterial cells to survive in free-living conditions where high anthropogenic Co^2+^ content may be encountered.

## Introduction

Cobalt (Co) and iron (Fe) are essential micronutrients in bacteria. While Fe is broadly required to deal with biological reactions in the actual conditions ruled by dioxygen, Co is involved in a handful of metabolic pathways as a catalytic cofactor or as the coenzyme vitamin B_12_ [1-3]. The chemical properties of both Fe and Co mean that they become toxic under certain conditions that promote increased bioavailability, i.e., high concentration in the extracellular environment, or in the case of Fe, an increase in the exchange between intracellular storage and the labile pool (*en route exchange*) [4, 5]. Although Fe toxicity seems to be curbed by Fe-storage systems, recent studies have suggested that Fe accumulation may exceed their buffering capacity and lead to redox stress [6, 7].

The rhizobial α-proteobacteria and alfalfa symbiont, *Sinorhizobium melliloti*, is involved in biological N_2_-fixation, a process highly dependent on Fe [8]. Previous studies describing genetic and mechanistic findings on Fe acquisition, export and storage in α-protobacteria, including rhizobia [9], share one of the two transcriptional factors, *irr* [10] and *rirA* [11], which sense cellular Fe status through heme or Fe-S clusters, respectively. RirA regulates iron uptake gene transcription, promoting free (ferric) and non-free (siderophore-iron complexes or heme) iron uptake in low Fe conditions. Irr regulates genes involved in heme synthesis and Fe exporters, derepressing their transcription in high Fe conditions. In *Agrobacterium tumefaciens* and *Bradyrhizobium japonicum*, the membrane-bound ferritin A (MbfA), a member of the CCC1 Fe efflux family, has been described to be involved in Fe homeostasis [12, 13]. In *S. meliloti*, a P-type-IB-ATPase (Nia) has also been described as being specific for Fe transport and upregulated under low O_2_ conditions [14]. Cobalt is an essential nutrient for *S. meliloti* and other rhizobia [15], and an uptake system has been identified in this organism [16]. Similarly, the *dmeRF* system, consisting of a member of the FrmR family of transcriptional regulators (DmeR) and a member of the cation diffusion facilitator (DmeF), has been involved in Co^2+^ homeostasis through export, specifically during free-living stages previous to establishing symbiosis and nodulation [17-19].

A feature of Co^2+^ dyshomeostasis is related to Fe-S cluster stability. There is evidence that a high Co^2+^ condition induces Fe-S cluster disassembly, triggering *suf* and *isc* machinery in response to this injury [20, 21]. The mechanisms proposed leading to Fe-S disruption by Co^2+^ or other transition metals vary from the non-cognate metal interfering with assembly process [20], to the Fe-S centers attacked in enzymes [22]. However, the final outcome in order to supply Fe for cluster reconstitution relies on changes in the dynamic between intracellular Fe stores and the labile-Fe pool (LIP) [23], and most importantly, the triggering of free and non-free Fe uptake mechanisms [18, 21, 24].

In rhizobia, the Fe-S cluster synthesis pathway is regulated by the transcription factor RirA, which represses the *suf* operon, the unique system for Fe-S cluster assembly in *S. meliloti* [9], under replete Fe conditions [11]. RirA is a functional analog of Fur, the canonical Fe-responsive transcriptional regulator, but differs from this in that it senses Fe indirectly through an Fe-S cluster with a labile Fe as a proxy [25, 26]. Thus, under low iron conditions, the release of RirA from the promoter region (IRO box) activates the transcription of the genes in its regulon. *S. meliloti* presents a second transcriptional regulator, Irr, which senses Fe status through heme binding and regulates *rirA* transcription, reinforcing the role of *rirA* under low iron conditions [4].

Some authors have proposed a link between Co^2+^ and Fe^2+^ homeostasis, providing evidence of a RirA-mediated response upon high extracellular Co^2+^, involving upregulation of heme uptake systems, probably as a result of an increase in the inactive apo form [18]. As a Co^2+^ Fe-S cluster attack could lead to an increase in Fe^2+^ bioavailability [27-29], an Irr-mediated response is also expected.

Importantly, previously described Co^2+^ transporters are emerging as Fe-exporters in bacterial organisms [7, 30, 31]. Therefore, we decided to evaluate the Fe transport capacity of SMc04167/AitP, a DmeF homolog in *S. meliloti*. Our data, showing that Co^2+^ induces Fe dyshomeostasis by likely affecting dynamics in the LIP, and that Co^2+^ sensitivity can be rescued in a non-competitive fashion by heme, together with the observations that AitP exports Fe^2+^, indicates a role for the FrmR/DmeR-regulated Co^2+^ exporters of the CDF family at initial stages of the Co^2+^ induced mediated Fe-S cluster salvage pathway through Fe^2+^ export.

## Results

### AitP is a classical CDF Co^2+^ exporter member of a sub-family with non-unique operon architectures

A survey of more than 17,000 complete bacterial genomes allowed us to identify by Best-Reciprocal-Hit more than 759 *dmeF*/*aitP*-like genes (Fig. 1A), showing typical features of this Co^2+^ CDF subfamily. A multigene blast analysis showed that approximately 62% of these share the archetypical operon organization of juxtaposed *frmR-aitP*/*dmeR-dmeF* loci (Fig. 1A). The remaining percentage appears to have an FrmR-independent regulation with no archetypical operon configuration and no operator-promoter region for the Co^2+^ responsive transcriptional regulator FrmR/DmeR (Suppl. Table 1). These findings also suggest that a significant number of putative Co^2+^ exporter *aitP*-like genes might be part of Co^2+^-responsive operons, with effectors other than Co^2+^ generated when this is accumulated. Moreover, AitP-like members found in Co^2+^-unresponsive operons could have a broaden specificity of transport.

**Figure 1.**
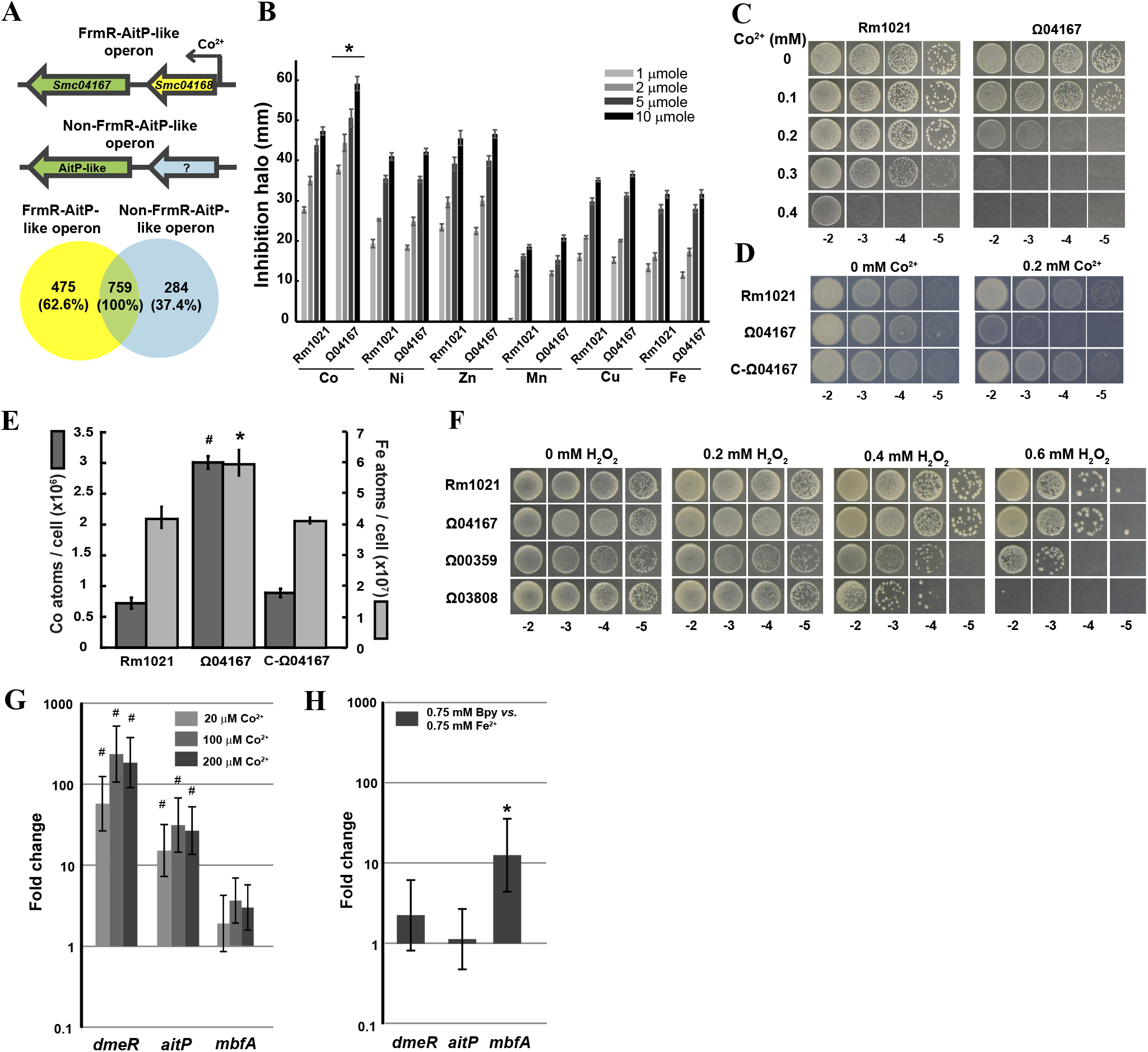
AitP is involved in Co^2+^/Fe^2+^ homeostasis and the genetic context of bacterial AitP-like transporters points to a broader transport specificity and regulation. A) Operon architecture analysis of AitP-like transporters in bacterial sequenced organisms indicates the existence of a subgroup with different transcriptional regulation. B) Metal toxicity was evaluated by inhibition halo assay using *S. meliloti* Rm1021 and Ω04167. Bars represent the inhibition halo diameter around 7-mm filter paper discs containing 10 (black), 5 (dark gray), 2 (gray) or 1 (light gray) μmole of the indicated metal (as Cl^-^ except for FeSO_4_). Notice the Co^2+^ sensitivity in Ω04167. * = p<0.01 *vs*. Rm1021. C) Serial dilutions (10 μl) of cells, initially at OD_600nm_=0.4, were spotted on TY-agar supplemented with the indicated Co^2+^ (as Cl^-^) concentration for 48 h at 30°C. D) Complementation was achieved by transformation of Ω04167 with pJB3Tc20-P_770_-*aitP* (C-Ω04167 strain). Notice that Co^2+^ sensitivity was restored to WT levels in the this strain. E) Cobalt (dark gray) or iron (light gray) content in cells after incubation with 0.2 mM CoCl_2_ for 1 h or 1.5 mM FeSO_4_ for 30 min at 30°C in TY medium during early exponential phase growth. Data are the mean ± SE of three independent experiments. * = *p*< 0.05 and # = *p*<0.01 from a two tail Student’s t-test *vs*. Rm1021. F) Serial dilutions (10 μl) of cells initially at OD_600nm_=0.4 were spotted on TY-agar supplemented with the indicated hydrogen peroxide concentrations for 48 h at 30°C. G) Fold change expression levels (2^-ΔΔCt^ with 16S as reference gene), of *dmeR, aitP* and *mbfA* with Co^2+^ or H) Fe^2+^ treatment with low (0.75 mM 2,2’-bipyridyl (Bpy)) or high Fe^2+^ (0.75 mM FeSO_4_). Cells were grown till mid-log phase at 30°C in TY. At this point media were supplemented accordingly and incubated for 45 min at 30°C before RNA extraction. Data are the mean ± SE of three independent experiments. * = *p* < 0.05 and # = *p*<0.01 from a two tail Student’s t-test of averaged ΔC_t_ (treated *vs*. non-treated/Bpy).

The genome of *S. meliloti* encodes a single member of the Co^2+^ efflux CDF subfamily (SMc04167, AitP), a homolog of the DmeF protein described in *R. leguminosarum* [17], and also of the AitP transporter of *Pseudomonas aeruginosa* [7]. As in other rhizobia, AitP is preceded by a transcriptional regulator of the FrmR/DmeR family. Phenotypic analysis of the insertional mutant of the SMc04167 locus in *S. meliloti* Rm1021 (Ω04167 or “AitP mutant”) shows increased Co^2+^ sensitivity in this strain (Fig. 1B). The sensitivity was concentration-dependent and it was rescued after gene complementation (Fig. 1C-D). Co^2+^ and Fe^2+^ accumulation occurred in the Ω04167 strain after incubation for 1 hour with 0.2 mM Co^2+^ or 20 minutes with 1.5 mM Fe^2+^,and complementation of Ω04167 reverted the accumulation to WT levels (Fig. 1E). Cell numbers after incubation were the same in all strains, indicating that these treatments (time and Co^2+^ or Fe^2+^ concentrations) are sub-lethal (not shown). Although the Fe^2+^ accumulation in AitP mutant did not induce a defect in cell growth, the double mutant lacking AitP and MbfA transporters was more sensitive to hydrogen peroxide (H_2_O_2_) than the MbfA single mutant in growth media with no external Fe added (Fig.1F). Free Fe^2+^ catalyzes the formation of reactive oxygen species through a Fenton reaction, which indicates that AitP functions as an Fe^2+^ exporter. As expected, transcription of *aitP* and *frmR*/*dmeR* was upregulated by Co^2+^(Fig. 1G) but was unresponsive to Fe^2+^ in the case of *aitP* as opposed to *mbfA* (Fig. 1H).

### AitP shows a dual specificity of transport

The phenotypical findings (Fig. 1E-F) agree with previous studies on AitP-like members [7], but suggest an alternative role in Fe^2+^ export by AitP. As Fe^2+^ homeostasis plays a significant role for the rhizobia-legume symbiosis, we decided to test whether Fe^2+^ could be transported by AitP. This could be hindered by the apparently redundant *mbfA* and *nia* Fe^2+^ transport systems [12-14]. Therefore, we heterologously overexpressed AitP in the Δ*fieF E. coli* strain (Suppl. Fig. 1A). FieF was previously described as an Fe^2+^ transporter [31, 32], and thus the genotypic background lacking FieF would allow measurement of Fe^2+^ with a low background signal. After protein induction, everted membrane vesicles were obtained and energized by pre-incubation with the electron donor NADH (or lactate), generating a H^+^-electrochemical gradient (Suppl. Fig. 1B). Only in this condition was ^59^Fe^2+^-importing activity to the vesicle detected, indicating that Fe^2+^ is a substrate transported by AitP, and that the driving force is the electrochemical gradient of H^+^ (Fig. 2A). After initial rate conditions were determined, we characterized the kinetics of the transport (Fig 2B). The estimated K_*1/2*_ value at the low millimolar range (1.35±0.21) may indicate a low affinity for transport, in line with the iron-saving strategies observed in almost all living organisms toward intracellular Fe^2+^. The H^+^-dependent transport at sub-saturating concentrations (Fe^2+^ = 2mM) was inhibited only by Co^2+^, indicating that competition at the transport site is possible (Fig. 2C). However, Co^2+^ accumulation observed in the AitP mutant (Fig. 1E) suggests a dual (Fe/Co) transport specificity, with both ions as cytosolic substrates, likely exported by a sequential mechanism involving binding to the same transport site. Considering the absence of Fe^2+^ regulation in *aitP* transcription, these results suggest a role for AitP in Fe^2+^ transport in conditions in which concomitant intracellular Co^2+^ accumulation occurs.

**Figure 2.**
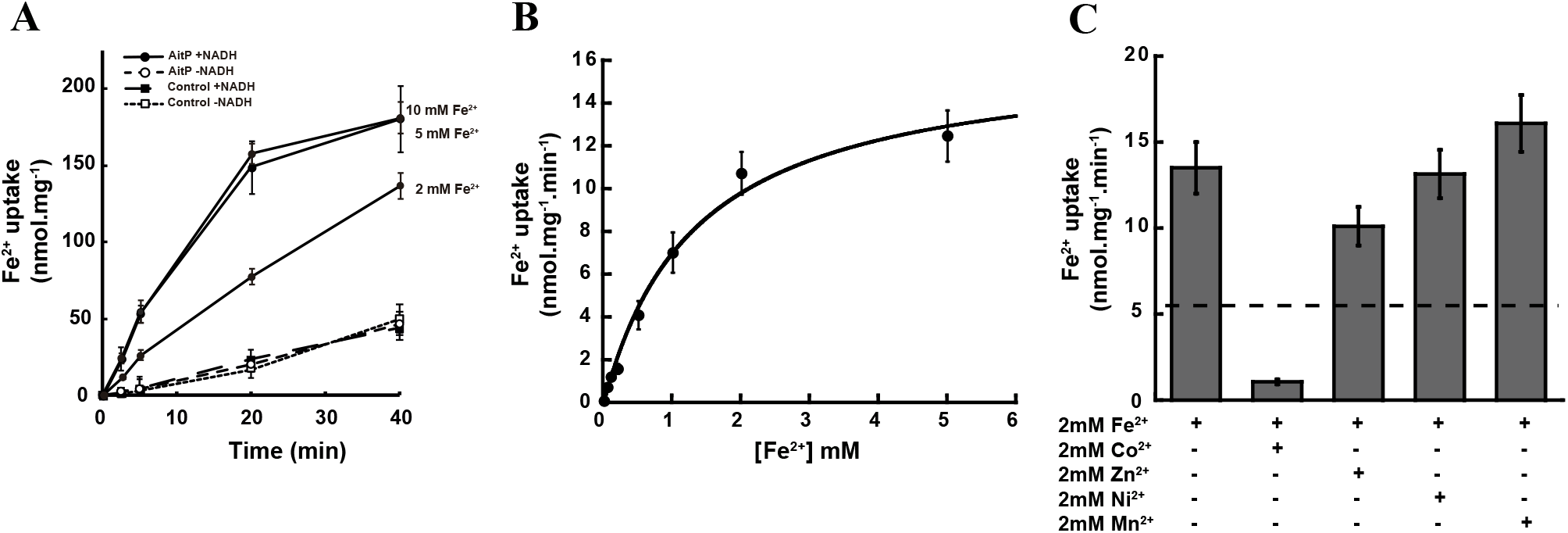
AitP transports Fe with low affinity and high specificity. Fe^2+^ uptake was measured in energized everted membrane vesicles (EMV) obtained from *E. coli* Δ*fieF* expressing AitP (see materials and methods). A) Time course and NADH dependency of Fe^2+^ uptake into EMV. Vesicles obtained from Δ*fieF* transformed with empty plasmid (empty and filled squares) or containing *aitP* (empty and filled circles) were assayed for NADH-dependent Fe^2+^ uptake activity. Notice that only AitP containing EMV (filled circles) present NADH-dependent activity. Initial rate conditions were assumed until 5 minutes after Fe^2+^ addition. Fe^2+^ concentrations for each trace are indicated (for control conditions only 5 mM traces are shown for clarity as the other concentrations did not show significant differences). B) Fe^2+^ transport kinetics. Data were fitted to *v*= V_max_ [Fe^2+^]/ K_*1/2*_ + [Fe^2+^] equation by using K_*1/2*_ = 1.35±0.21 mM and V_max_ = 16.4±1.0 nmol.mg^-1^.min^-1^. C) Apparent effects on Fe^2+^ transport rate by AitP of unlabeled divalent transition metals (TM^2+^) as real substrates or competitive inhibitors. Dashed line depicts the expected transport rate value obtained using *v*= V_max_ [Fe^2+^]/ (K_*1/2*_ (1+ ([TM^2+^]/K_i_))+[Fe^2+^]), with [Fe^2+^]=[TM^2+^]= 2 mM and K_i_ assumed to be equal to K_*1/2*_. Note that only Co^2+^ affects the Fe^2+^ transport rate negatively, probably as real substrate. In all cases, Fe^2+^ uptake by vesicles from empty vector-transformed strain *E. coli* Δ*fieF* was subtracted prior to data analysis. In all cases, data are the mean ± SE of three independent experiments.

### AitP participates in Fe-homeostasis

Previous phenotypical characterizations of *aitP*-like genes in other rhizobial organisms point to a similar role of AitP in Co^2+^ export and detoxification [17-19], and we have previously shown that the AitP homolog in *P. aeruginosa* is involved in Fe^2+^ homeostasis [7]. However, a P-IB-type ATPase, *nia*, has been described as participating in Fe^2+^ homeostasis in this organism [14], and the identification of an *mbfA* ortholog in Rm1021 genome led us to hypothesize that AitP in *S. meliloti* may play a non-classical role in Fe^2+^ homeostasis. In this line, Fe^2+^ sensitivity was not observed in the Ω04167 strain, pointing to *nia* and *mbfA* as major exit routes for Fe^2+^ in rhizobia. As Co^2+^ transport has been described through Fe^2+^ exporters, we conducted phenotypical analyses to test whether double mutants showed increased sensitivity towards Co^2+^. We observed an increased sensitivity in the double mutants, Ω3004 and Ω03808, beyond the threshold shown by the Ω04167 strain (Fig. 3A) but, interestingly, no significant increases of Co^2+^ accumulation were observed *vs*. the Ω04167 strain (Fig. 3B). Supporting the exclusive participation of AitP in Co^2+^ efflux, Co^2+^-dependent *nia* transcription is not expected [14], and *mbfA* was not induced by Co^2+^ (Fig. 1G). Thus, we hypothesized a possible Fe^2+^-induced toxic effect being added to those promoted by Co^2+^ accumulation. To test this, we measured the Co^2+^ sensitivity of the Ω04167 *vs*. Rm1021 in the presence of Fe. The increased sensitivity shown by Ω04167 in this condition (Fig. 3C) indicates that Co^2+^ accumulation induces a shift in iron distribution, with increased LIP or higher levels of Fe^2+^ bioavailability that result toxic to the cell. As the LIP can be fed with intracellular and extracellular Fe sources, we compared the transcription of *fbpA*, the substrate binding protein of the ferric cation ABC transporter, in WT and AitP mutant exposed to sub-lethal Co^2+^ concentrations, as a proxy of extracellular Fe contribution. We detected a significant increase of *fbpA* in both strains (Fig. 3D), with higher levels in the AitP mutant, indicating that Fe import and probably intracellular Fe contribute to Co^2+^ toxicity. Considering that the Co^2+^ sensitivity of double mutants was observed in media with no added Fe (ca. 10 μM total Fe), we tested the effect of Co^2+^ on pathways whose inhibition could lead to an increase in the bioavailability of iron. As Co^2+^ inhibits heme synthesis and induces Fe-S disassembly, thus promoting the increase of LIP, we quantified the total protoporphyrin content in both strains, exposed or not to sub-lethal Co^2+^ (Fig. 3E) and measured the transcription of *sufA* in WT, a gene upregulated in response to Fe-S cluster attack. Although Co^2+^ treatment decreases the level of protoporphyrin, this is similar in both strains, suggesting that the Co^2+^-mediated Fe-S cluster attack is the main contributing source to LIP. Regarding this, we observed *sufA* up-regulation in WT in the presence of sub-lethal Co^2+^ concentrations (Fig. 3F). We also analyzed the Co^2+^-dependency of transcriptional levels of bacterioferritin (*bfr*) and bacterioferrodoxin (*bfd*) [33], as an estimate of the Fe storage status and the RirA-mediated response. Only *bfd* transcription was increased by Co^2+^ (Fig. 3F). Finally, Fe-S cluster or heme synthesis disruption would lead to increased Fe^2+^ bioavailability, and a suitable proxy to test this are the *mbfA* and *bfr* transcriptional levels, but increases were not statistically significant (Fig. 1G and 3F).

**Figure 3.**
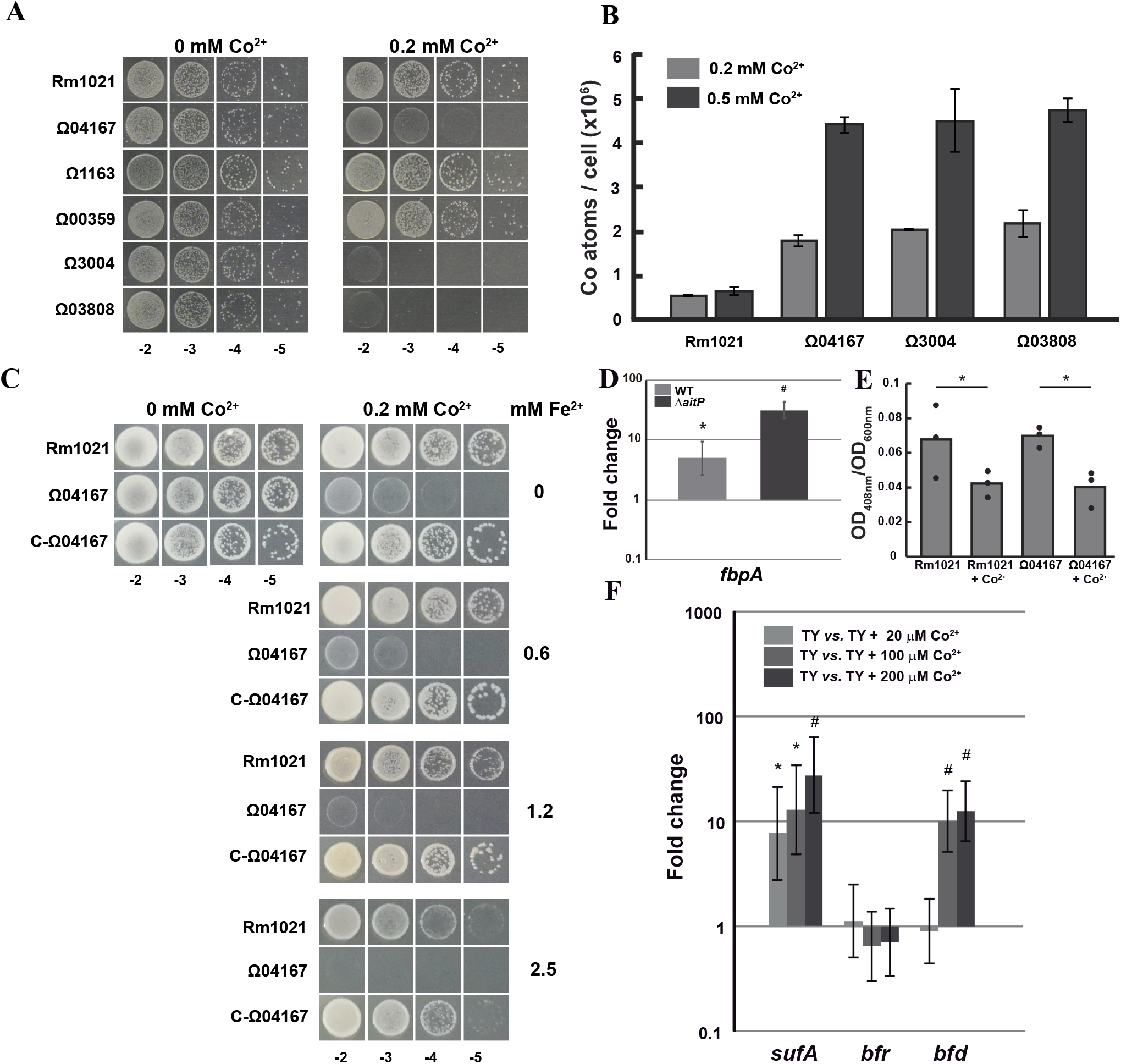
AitP is involved in Fe^2+^ homeostasis. A) Serial dilutions (10 μl) of *S. meliloti* Rm1021, Ω04167, Ω1163, Ω00359, Ω3004, and Ω03808 cells, initially at OD_600nm_=0.4, were spotted on TY-agar supplemented with the indicated Co^2+^ (as Cl^-^) concentration for 48 h at 30°C. B) Cobalt content in Rm1021, Ω04167, Ω3004, and Ω03808 after incubation with 0.2 or 0.5 mM Co^2+^ for 1 hour at 30°C in TY medium during early exponential phase growth. C) Serial dilutions (10 μl) of *S. meliloti* Rm1021, Ω04167, and C-Ω04167 cells, initially at OD_600nm_=0.4, were spotted on TY-agar supplemented with the indicated Co^2+^ (as Cl^-^) and Fe^2+^ (as SO_4_^2-^) concentration for 48 h at 30°C. D) Fold change expression levels (2^-ΔΔCt^ with 16S as reference gene) of *fbpA*. Cells were grown till mid-log phase at 30°C in TY. At this point media were supplemented with 200 μM CoCl_2_ and incubated for 45 min at 30°C before RNA extraction E) Total heme was estimated after cell incubation with 0.2 mM CoCl_2_ for 2 hours at 30°C during the exponential growth phase. Data points and the mean of three independent experiments are shown. * = *p*<0.05. F) Fold change expression levels (2^-ΔΔCt^ with 16S as reference gene) of *sufA*. Cells were grown till mid-log phase at 30°C in TY. At this point media were supplemented accordingly and incubated for 45 min at 30°C before RNA extraction. For D) and F) Data are the mean ± SE of three independent experiments. * = *p*<0.05 and # = *p*<0.01 from a two tail Student’s t-test of averaged ΔC_t_ (treated *vs*. non-treated).

### AitP Fe^2+^ transport likely orchestrates the initial stages of the Co^2+^-induced Fe-S cluster salvage pathway

As Co^2+^-dependent Fe^2+^ toxic effects seem mainly related to Fe-S clusters, the AitP function may be related to driving a concerted RirA-dependent Fe-S cluster salvage/synthesis pathway. If Co^2+^ accumulation led to Fe-S disassembly, we hypothesized that genes regulated by RirA might be dysregulated, similarly to that observed for *fbpA* and *sufA* (Fig 3D and F). In particular, SMc02726 or *shmR*, a hemin-binding outer membrane receptor [34], is induced in low Fe conditions and in a Δ*rirA* mutant [35]. We detected increased transcripts of the outer membrane porin, ShmR, and the periplasmic component, HmuT, of the ABC-heme importer HmuTUV [36](Fig. 4A). This upregulation of the heme uptake system in the Ω04167 mutant suggested an exacerbated response to Co^2+^ accumulation, likely due to an induced shift in the equilibrium of RirA towards the apo-form. We then tested if adding hemin in the media affected the Co^2+^ sensitive phenotype detected in the Ω04167 strain. As shown in Fig. 4B, hemin addition to buffered TY medium (pH = 6,5) with 0.2 mM Co^2+^ rescued the Co^2+^-sensitive phenotype at concentrations between 50-200 μM. Chelation of Co^2+^ by Fe-protoporphyrin IX (hemin) or metal exchange was discarded, as empty protoporphyrin IX (PPIX) at 200 μM did not decrease Co^2+^ sensitivity (Fig. 4C). The detection of increased transcripts of hemin transport systems in Co^2+^ treated cells led us to test the hemin uptake rate under the same treatment. Surprisingly, the AitP mutant did accumulate hemin at a significantly lower rate *vs*. WT (Fig. 4D and E). Although counterintuitive, this result indicates that an Fe/Co-dependent post-transcriptional regulation is possible. Overall, the data point to a role for AitP in Fe export under Co-rich conditions in order to concert a RirA-dependent heme uptake process likely used for Fe-S cluster assembly.

**Figure 4.**
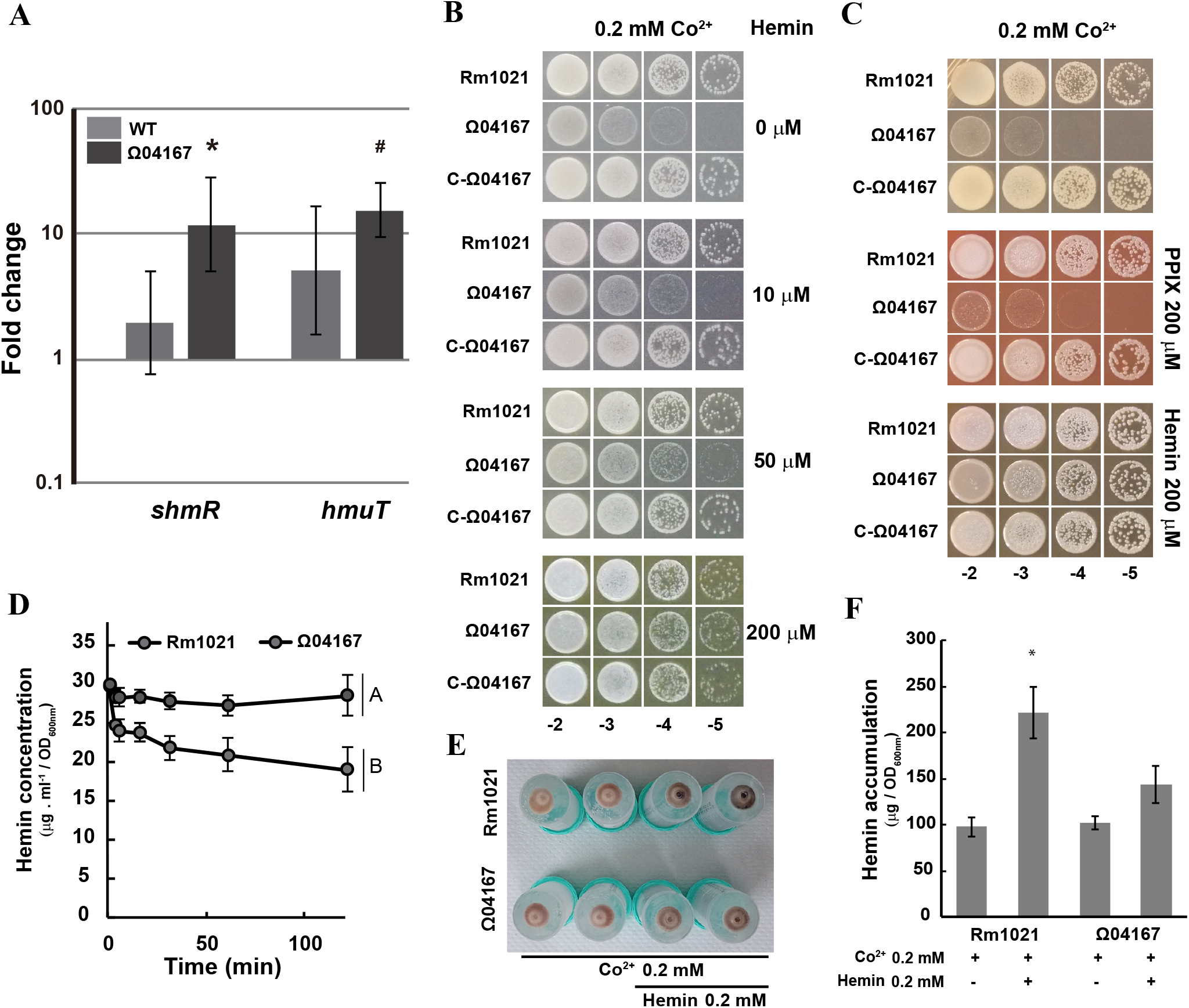
AitP activity induces heme uptake transport. A) Fold change expression levels (2^-ΔΔCt^ with 16S as reference gene) of heme uptake components in WT (light gray) and Ω04167 (dark gray) *in vitro vs*. 16S reference gene. Cells were grown till mid-log phase at 30°C in TY. After this, half of the culture volume was supplemented with CoCl_2_ or not, and incubated for 2 hours at 30°C before RNA extraction. Data are the mean ± SE of four independent experiments. * = *p*<0.05 and # = *p*<0.01 from a two-tail Student’s t-test of averaged ΔC_t_ (treated *vs*. non-treated). B) Serial dilution of *S. meliloti* Rm1021, Ω04167, and C-Ω04167 cells, initially at OD_600nm_=0.4, were spotted (10 μl) on TY-agar supplemented with the indicated Co^2+^ (as Cl^-^) and hemin concentration for 48 h at 30°C. C) Serial dilutions of Rm1021, Ω04167, and C-Ω04167 cells, initially at OD_600nm_=0.4, were spotted (10 μl) on TY-agar supplemented with the indicated Co^2+^ (as Cl^-^) and 200 μM PPIX or 200 μM hemin concentration for 48 h at 30°C. D) Hemin uptake assay of WT (light gray) and Ω04167 (dark gray) cells. Different letters designate significantly different means as informed by a Bonferroni post hoc (p<0.05) ANOVA test. E) Illustrative photograph of cell pellets showing Co^2+^ induced hemin uptake. Cells were grown in TY at 30°C till mid-exponential phase and then incubated with Co^2+^ and hemin concentrations as indicated for 2 hours (n=2). After several washes, pellets were imaged. Notice the darker green color of Rm1021 cells (upper-right) after the Co^2+^+hemin incubation *vs*. Δ*aitP*, indicating increased hemin accumulation. Picture shows representative results of at least three independent experiments. F) Quantification of total hemin in cell pellets. *=p<0.01 vs Rm1021.

## Discussion

Cobalt is an essential element for life, and organisms like *S. meliloti* have a special requirement for Co for methionine synthesis [15, 16]. However, in high concentrations it proves toxic to cells. Co^2+^ efflux transporters of the CDF family have been previously characterized in rhizobia, and their transcriptional regulation is mainly governed by a FrmR/DmeR transcription factor [17]. Our bioinformatics analyses show that a significant number of members of this family may be regulated by different factors, thus responding to other effectors. Interestingly, several of these members appear under the regulation of the Multiple Antibiotic Resistance Regulator (MarR) superfamily of transcription factors. Although analysis of the functional roles of non-FrmR-regulated Co^2+^-CDF is beyond the scope of this work, the heterogeneity in *aitP*-like homolog regulation points to their capacity to transport other divalent cations. Recent works have expanded the substrate of the Co^2+^ efflux CDF transporters [7], and in this work, we have unequivocally confirmed the Fe^2+^ transporting capacity of AitP by ^59^Fe^2+^ transport assays in energized everted vesicles. Due to the sensitivity of the mutant to high Co^2+^ but not to Fe^2+^, from a physiological perspective two questions arise: why does AitP export iron? And why is it regulated by Co^2+^?

The main toxic effect of Co^2+^ in bacteria seems related to Fe-S cluster assembly [20]. In *E. coli*, the responses to the intoxication involve the induction of the Fe-S cluster salvage pathway in order to counterbalance this effect, but also, downregulation of free-Fe import systems, such as Feo [23]. In contrast, *S. meliloti*, and likely all α-proteobacteria of the *Rhizobiaceae* clade bearing RirA homologs, handle Co^2+^ toxicity by inducing iron uptake mechanisms, i.e., triggering an Fe-starvation-like response. In *Agrobacterium tumefaciens*, RirA represses genes related to free and non-free Fe uptake, and the repression is lost in the presence of Co^2+^ [18]. Our results also point to an induction of the expression of these iron uptake systems by Co^2+^ (*fbpA, shmR* and *hmuT*; Fig 3D and 4A). Similarly, this is prompted by a derepression involving Co^2+^ attack on holo-RirA.

It has been proposed that the Fe-S cluster attack might lead to an increase of the labile-Fe pool (LIP). In the presence of Co^2+^ intoxication and accumulation, the contribution of Fe from the Fe-S cluster to the LIP may be detrimental by oxidative damage. We detected *suf* up-regulation induced by Co^2+^, which is in line with the increase. This could be an important Fe source in low iron conditions, likely contributing to the increase in sensitivity observed in double mutants lacking the main Co^2+^ efflux system and one of the Fe^2+^ exporting systems (Fig 3A). As AitP-like members are also found in δ-proteobacteria, which mostly regulate Fe homeostasis via direct sensing by Fur, Fe^2+^ export activity may be required to promote apo-Fur and the Fe-starvation response through non-free Fe import.

Thus, AitP activity counterbalances the increase of LIP coming both from the release of Fe from Fe-S clusters disassembled by Co^2+^ and from the Fbp-mediated entry. The latter, although it represents an apparently futile Fe import-export cycle, is promoted only by Co^2+^ accumulation. In agreement with our hypothesis, the addition of free iron increased Co^2+^ toxicity in the AitP mutant (Fig. 3C), while the non-free iron from hemin had a protective role (Fig. 4B,C), likely allowing replenishment of the intracellular pool without incurring in stress for the bacteria.

Two more plausible, but still speculative mechanisms, involve RirA and the regulatory protein, *hmuP*/*hemp* [37]. Direct attack on RirA by Co^2+^ explains the transcriptional derepression observed of Fe-S cluster salvage (Fig. 3F) and hemin uptake genes (Fig 4A), but still leaves the export role of Fe for AitP dispensable. A role in the fine tuning of the Co^2+^ toxic response involving Fe-export by AitP might be explained considering the post-transcriptional regulation of *shmR* in rhizobia [38]. A unique putative binding site for the HmuP protein (a regulatory protein of *shmR*) was found in 5’UTR of *shmR* transcripts. Importantly, heme binding to HmuP has been shown *in vitro* [39]. We speculate that *in vivo* HmuP sensing capacity, likely via free-Fe or PPIX binding, may be involved in the post-transcriptional regulation of *shmR*. Thus, a LIP increase in the AitP mutant would downregulate *shmR* expression. Our data showing lower hemin uptake capacity of AitP mutant *vs*. WT, even when transcripts are elevated, support this. The recovery of Co^2+^ resistance in the presence of high hemin concentrations (Fig. 4B) points to an unspecific uptake, or passive diffusion, mechanism in the AitP mutant. These hypotheses are currently being tested.

Based on the above, we propose a model during Co^2+^ dyshomeostasis in which AitP would detoxify both Co^2+^ and Fe^2+^. This dual export allows cells to reduce the primary stressor (Co^2+^), while maintaining the labile-Fe pool in range despite the release of free Fe from the Fe-S clusters or the entry of free Fe via ABC transporter. The lack of heme import activity, even when transcripts of the importing systems are high, suggests that AitP activity is involved in the orchestration of the transcription-translation process of ShmR and Hmu. Future studies will focus on the role of AitP-like members in the regulation of heme uptake systems and the need for other partners to achieve this regulation.

## Materials and methods

### Bioinformatic analyses

To analyze the relative abundance of the canonical “FrmR-AitP” operon architecture, *S. meliloti* AitP (SMc04167/ SM_RS10235) homologs were searched by the best reciprocal hit method [40]. Briefly, a database was created locally with all the protein sequences from bacteria with a complete genome (17,972) and chromosome (2,570) assembly level obtained from ftp://ftp.ncbi.nih.gov/genomes/refseq/bacteria/assembly_summary.txt (August 2021). A BLAST search in this database, with WP_010969625.1 (SMc04167/ SM_RS10235) as query and an Expect-value of 1.10^−5^, resulted in 13,570 subject/hits. Removal of duplicated subject/hits by accession number led to 4,431 SMc04167 homolog candidates. These were blasted against *S. meliloti* 1021 (GCF_002197445.1_ASM219744v1_protein.faa) individually. In all cases BLAST retrieved WP_010969625.1 as the subject, confirming best-reciprocal-hits association. After visual inspection of the subject annotations, CDFs lacking the canonical poly-histidine stretch sequence (TMS4 and TMS5) were excluded in order to minimize the number of arbitrarily defined “true candidates” (Expect-value<2.19.10^−35^ and percentage of positive-scoring matches>54%). There were 1,651 assemblies of bacterial genomes having AitP-like members with unique accession numbers. This number of organisms reflects several strain assemblies of the same organism, and we maintained it in order to promote the sensitivity of the final assessment. These were used later to compare and create a list of 759 non-redundant bacterial organisms having, or not having, the FrmR-AitP architecture. The FrmR-AitP architecture search was performed using Multigenblast [41]. For these analyses, the genomic database was built with approximately 4,200 genebank-formatted (.gbbf) assemblies of chromosome and complete genome with identical gender and species names to those drawn from the 1,651 organisms within the AitP list. The SMc04168-SMc04167 sequence was used to run the analyses. Results were parsed to obtain the final list of organism names with the operon architecture (∼500) and compared to the list of the AitP-possessing organisms (∼800).

The remaining organisms with non-FrmR-AitP-like architecture (∼300) were further analyzed. Three search strategies were followed to confirm the lack of FrmR(RcnR/DmeR)-dependent regulation in those organisms with AitP-like members lacking the FrmR-AitP architecture. In the first, the 500bp up-stream CDS region was downloaded and used in a tblastn search, using the protein sequence of SMc04168/*dmeR* as query. In the second, protein accession numbers of AitP-like members of this group were used to retrieve the up- and downstream loci annotations, using the E-fetch “identical protein group” tool. A third strategy consisted in performing a blastn search in the 500bp upstream regions for the consensus FrmR consensus binding motif, using the *dmeR* binding sequence up-stream SMc04167 (AAATAGGGTACCCCCCTATGCT) as query. These searches allowed us to identify 27 non-FrmR-AitP-like previous candidates as false negatives, and to confirm that 284 AitP-like loci lack the FrmR-AitP architecture. After removal of duplicated organism names, a group diagram was prepared with these numbers.

### Culture media, strains and mutant constructions

*S. meliloti* strains (Table 1) were cultured in TY media (0.5% tryptone, 0.3% yeast extract, 0.09% CaCl_2_) supplemented with streptomycin 600 μg/ml. The *S. meliloti* strain Rm1021(StrR) and *E. coli* S17-1 [42] carrying the plasmid pK19mob2ΩHMB for the mutation of SMc04167 were obtained from University of Freiburg, Germany (Table 1). The deletion mutant strain of gene SMc04167 (Ω04167) was generated by plasmid insertion following described protocols [43]. Briefly, freshly grown *S. meliloti* Rm1021 was mixed with *E. coli* S-17 cells bearing the plasmid pK19mobΩHMB, containing the internal sequence of the gene SMc04167, named S17-1.PI.G1PELR20F8 (www.rhizogate.de (GenDB)), and then incubated overnight at 30°C on a nitrocellulose filter disk placed on LB-agar. Transconjugants were recovered from the filter and selected by plating on TY-agar supplemented with streptomycin 600 μg/ml and kanamycin 50–200 μg/ml. Complementation was achieved by conjugation of Ω04167 with the non-integrative plasmid pJB3Tc20-P_770_-*aitP* containing gene SMc04167 plus 770 bp upstream. Transconjugants were selected on TY media with kanamycin (200 μg/ml) and tetracycline (10 μg/ml). Single and double mutants of *nia* (SMa1163) and *mbfA* (SMc00359) were generated by conjugation and single crossover of plasmid pJQ200SK, containing a 600 bp internal region of each loci in Rm1021 (single mutant) or Ω04167 (double mutants). Transconjugants were selected in TY agar supplemented with streptomycin 600 μg/ml and gentamycin (50 μg/ml). Mutation of *fief* was performed as already described[44]. Briefly, 50 bp up- and downstream were joined to the Kan^R^ cassette from plasmid pKD4. After purification, the fragment was introduced by electroporation (1.8 kV, 600 Ω and 10 μF, 1mm gap cuvette) into BL21(DE3) cells previously transformed with pKD46. In order to induce the Rec system and to increase recombination, the electrocompetent cells were prepared by growing cells in LB supplemented with 1mM L- (+)-arabinose at 30°C. After electroporation, cells were recovered in 5 ml of SOC media. Transformants were selected in LB-agar plates with 50 μg/ml kanamycin incubated at 37°C for 16 hours. Curation of the Red system was confirmed in isolated colonies that showed negative growth after 16 h incubation at 37°C in LB-agar plates supplemented with 100 μg/ml ampicillin and 50 μg/ml kanamycin, but positive growth in plates supplemented with 50 μg/ml kanamycin. All strains were checked by PCR using purified genomic DNA as a template and internal and external primers (Table 1).

**Table 1.**
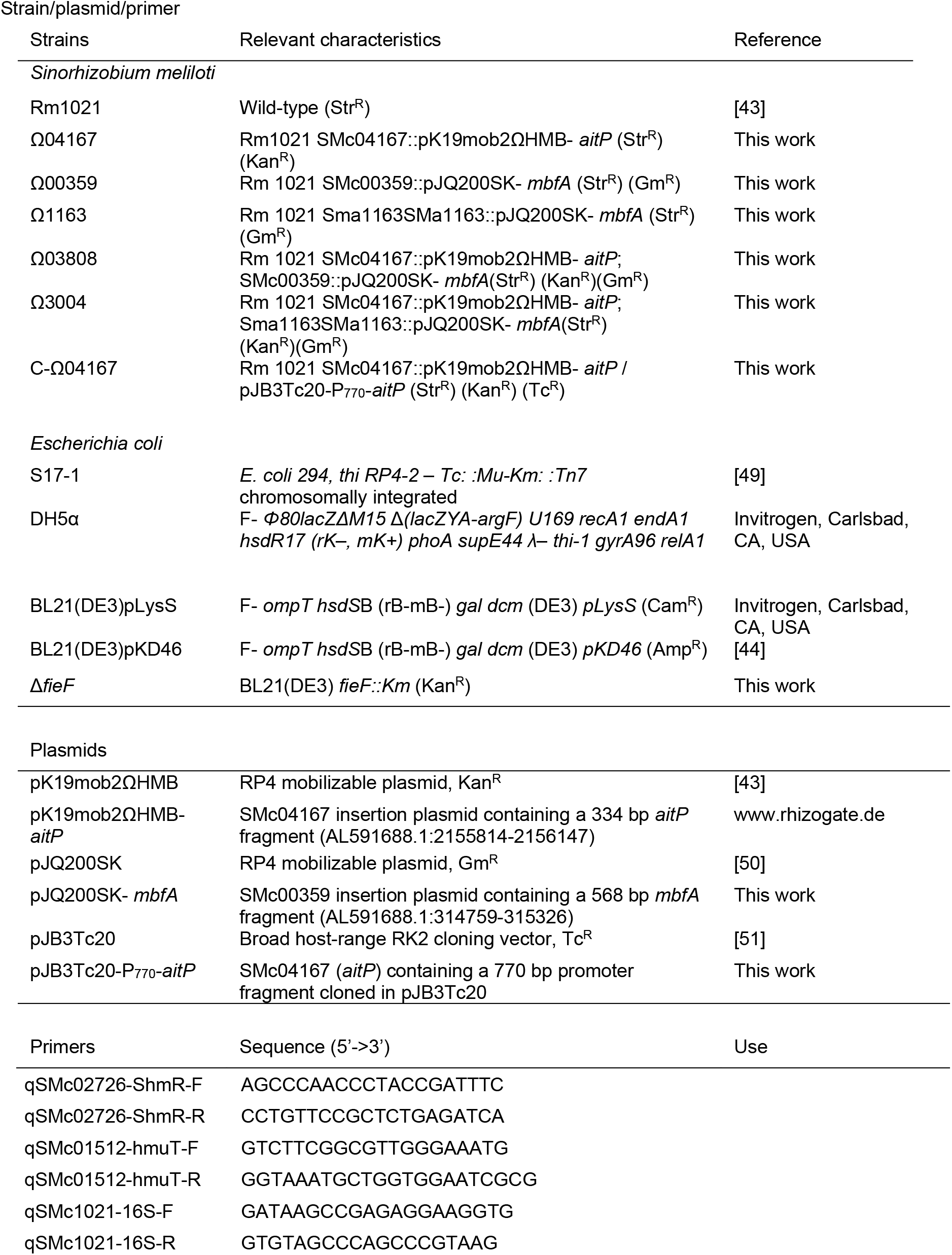
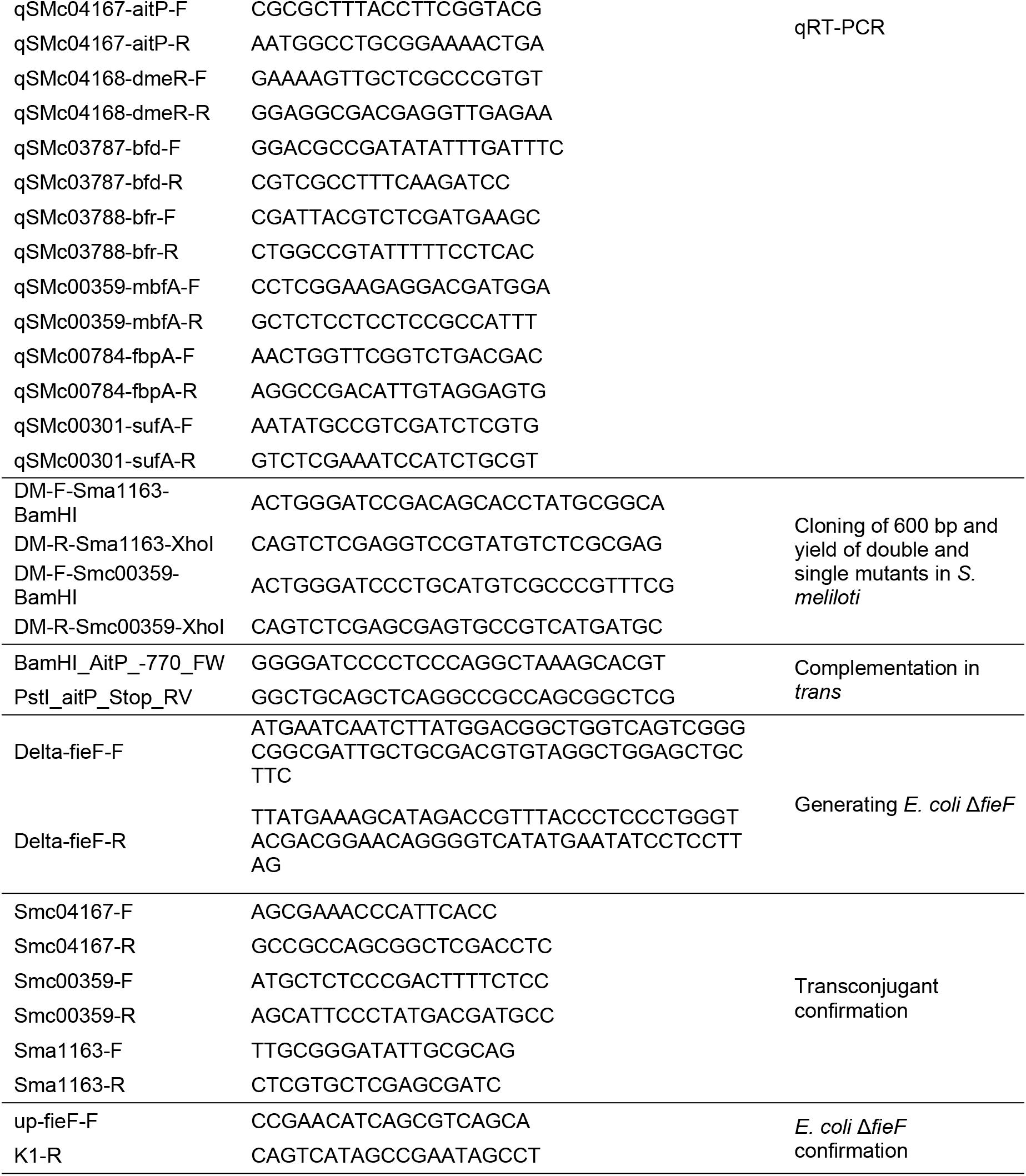
List of strains, plasmids and primers used in this study.

### Sensitivity tests

Serial dilutions of *S. meliloti* Rm1021, Ω04167, Ω00359, Ω1163, Ω03808 and Ω3004 cells, initially at OD_600nm_=0.4, were spotted (10 μl) on TY-agar supplemented with the indicated transition metals (as chloride salts, except Fe^2+^ as SO_4_^2-^)^-^ or H_2_ O_2_ concentrations for 48 h at 30°C. When hemin or PPIX (dissolved in 50 mM NaOH) were added to the media, the pH was buffered with MES-KOH 15 mM pH 6.5, and the control conditions (0 mM hemin, 0 mM PPIX) were supplemented with the same amounts of NaOH.

### Cobalt and iron accumulation assays

Cells were incubated with 0.1 mM CoCl_2_ for 1 hour or with 1.5 mM FeSO_4_ in TY medium during early exponential phase growth. Cells were pelleted and washed with 0.9% NaCl and mineralized with HNO_3_. Co or Fe were quantified by ICP-MS or Atomic absorption spectroscopy.

### Transcriptional analyses by qRT-PCR

Fold change expression levels (2^-ΔΔCt^) of iron exporters and iron-related genes were assayed under 20 μM, 100 μM or 200 μM CoCl_2_, and low (0.75 mM 2,2’-bipyridyl (Bpy)), or high (0.75 mM FeSO_4_) Fe^2+^ conditions *in vitro vs*. 16S reference gene. Cells were grown till the mid-log phase at 30°C in TY. After this, the culture volume was split, supplemented or not, and incubated for 45 min at 30°C before RNA extraction. Primers used are described in Table 1.

### Iron transport assay

Fe^2+^ uptake was measured in energized everted membrane vesicles (EMV) obtained from *E. coli* Δ*fieF* expressing AitP. Heterologous expression was detected by electroblotting SDS-PAGE Tris-glycine gels onto nitrocellulose membranes and immunostaining with primary rabbit anti-His-tag antibody and secondary goat anti-rabbit IgG phosphatase alkaline conjugated antibody (GenScript, Piscataway, NJ, USA) (Suppl. Fig. 1). Vesicles were obtained as previously described [45, 46] Before transport measurement, 10 μl vesicles (or the equivalent to 100 μg total protein of EMV) were energized in 100 μl of transport buffer (100 mM HEPES-Na, 250 mM sucrose, 50 mM KCl, 2 mM MgCl_2_, 5 mM TCEP, pH 6.8) by 2-minute pre-incubation with 2.5 mM NADH at 37°C and the uptake was initiated by addition of a 10-fold concentrated ^59^Fe^2+^ solution and mixing. Reaction was stopped after 5-minute incubation at the same temperature by filtering through 0.22 μm nitrocellulose. Filters were immediately washed with a standard solution containing 2 mM EDTA and dissolved in scintillation cocktail for counting.

### Hemin uptake assay

Hemin uptake was estimated following a previously described protocol [47]. Briefly, stationary phase cells cultured in 50 ml of TY medium were incubated for 2 hours with 0.2mM CoCl_2_. Cells were centrifuged, washed once with ice-cold phosphate-buffered saline (PBS) pH 7.45 and resuspended in 50 ml PBS 0.05 mM CoCl_2_ supplemented with 30 ug/ml of a hemin solution. At 2.5, 5, 15, 30, 60 and 120 minutes, one milliliter of culture was centrifuged and the supernatant collected to measure the absorbance at 400 nm. No significant difference was observed in the cell numbers of WT and AitP mutant strains after treatment, indicating that the incubation time with Co^2+^ used does not affect cell viability.

### Porphyrin quantification

Porphyrins were quantified following the described protocol [48]. Briefly, mid-log phase cells cultured in 50 ml of TY medium were incubated for 2 hours with or without 0.2mM CoCl_2_. Cells were centrifuged, washed twice with ice-cold 50 mM Tris-HCl (pH 8.0), resuspended in ethyl acetate/glacial acetic acid (3:1, v/v) to 1/100 the original culture volume, and lysed by sonication on ice. Cell debris was removed by centrifugation at room temperature, and the non-aqueous (top) phase was washed twice with 1 ml ddH_2_O to remove residual water-soluble contaminants. Extraction of porphyrins from the solution was achieved by the addition of 0.5 ml 3 M HCl, and the absorbance of the bottom-aqueous phase was assessed at 408 nm. Porphyrin levels were normalized to optical cell density (OD_600nm_). No significant difference was observed in the cell numbers of WT and AitP mutant strains after treatment, indicating that the incubation time with Co^2+^ used does not affect cell viability.

## Author contributions

Conceived and designed the experiments: D.R., M.G.-G, I.A., P.M., T. M.

Analyzed the data: D.R., M.G.-G, I.A., P.M., T. M.

Contributed reagents/samples/analysis/tools: D.R., M.G.-G, I.A., P.M., T. M.

Wrote or edited the manuscript: D.R., M.G.-G, I.A., P.M., T. M.

All authors read and approved the final manuscript.

## Abbreviations

TM: transition metal
CDF: cation diffusion facilitator
TMS: transmembrane segment;
LIP: labile-Fe pool
ICP-MS: inductively coupled plasma mass spectrometry
Bpy: 2,2’-bipyridyl.

## Acknowledgments

We thank Dr. J.M Argüello for help and support on the setting of ^59^Fe^2+^ transport assays.

