## Supplemental Table 1 for "Functional characterization of the Co^2+^ transporter AitP in *Sinorhizobium meliloti*: a new player in Fe^2+^ homeostasis"

| Bacterial organism with AitP-like | Bacterial organism with FrmR-AitP-like | Bacterial organism with non-FrmR-AitP-like |
|---|---|---|
| Achromobacter denitrificans | Achromobacter denitrificans | Acaryochloris marina |
| Achromobacter sp. MFA1 R4 | Achromobacter sp. MFA1 R4 | Acidihalobacter prosperus |
| Achromobacter spanius | Achromobacter spanius | Acidovorax ebreus |
| Achromobacter xylosoxidans | Achromobacter xylosoxidans | Acidovorax sp. KKS102 |
| Achromobacter | Achromobacter | Acinetobacter calcoaceticus/baumannii complex |
| Acidibrevibacterium fodinaquatile | Acidibrevibacterium fodinaquatile | Acinetobacter calcoaceticus |
| Acidiphilium cryptum | Acidiphilium cryptum | Acinetobacter guillouiae |
| Acidiphilium multivorum | Acidiphilium multivorum | Acinetobacter oleivorans |
| Acidovorax sp. 1608163 | Acidovorax sp. 1608163 | Advenella kashmirensis |
| Acidovorax sp. JS42 | Acidovorax sp. JS42 | Algicoccus marinus |
| Acinetobacter baumannii | Acinetobacter baumannii | Altererythrobacter atlanticus |
| Acinetobacter bereziniae | Acinetobacter bereziniae | Altererythrobacter sp. B11 |
| Acinetobacter defluvii | Acinetobacter defluvii | Anabaena cylindrica |
| Acinetobacter dispersus | Acinetobacter dispersus | Anaeromyxobacter dehalogenans |
| Acinetobacter gyllenbergii | Acinetobacter gyllenbergii | Anaeromyxobacter sp. Fw109-5 |
| Acinetobacter haemolyticus | Acinetobacter haemolyticus | Anaeromyxobacter sp. K |
| Acinetobacter indicus | Acinetobacter indicus | Arcobacter cibarius |
| Acinetobacter johnsonii | Acinetobacter johnsonii | Arcobacter nitrofigilis |
| Acinetobacter junii | Acinetobacter junii | Azoarcus sp. CIB |
| Acinetobacter lactucae | Acinetobacter lactucae | Azospirillum |
| Acinetobacter larvae | Acinetobacter larvae | Azotobacter chroococcum |
| Acinetobacter nosocomialis | Acinetobacter nosocomialis | Azotobacter salinestris |
| Acinetobacter pittii PHEA-2 | Acinetobacter pittii PHEA-2 | Beggiatoa leptomitoformis |
| Acinetobacter pittii | Acinetobacter pittii | Betaproteobacteria bacterium GR16-43 |
| Acinetobacter radioresistens | Acinetobacter radioresistens | Blastochloris tepida |
| Acinetobacter schindleri | Acinetobacter schindleri | Blastomonas fulva |
| Acinetobacter shaoyimingii | Acinetobacter shaoyimingii | Blastomonas sp. RAC04 |
| Acinetobacter sp. ACNIH1 | Acinetobacter sp. ACNIH1 | Bradyrhizobium amphicarpaeae |
| Acinetobacter sp. ACNIH2 | Acinetobacter sp. ACNIH2 | Bradyrhizobium betae |
| Acinetobacter sp. C16S1 | Acinetobacter sp. C16S1 | Bradyrhizobium canariense |
| Acinetobacter sp. LoGeW2-3 | Acinetobacter sp. LoGeW2-3 | Bradyrhizobium cosmicum |
| Acinetobacter sp. Marseille-Q1620 | Acinetobacter sp. Marseille-Q1620 | Bradyrhizobium diazoefficiens |
| Acinetobacter sp. MYb10 | Acinetobacter sp. MYb10 | Bradyrhizobium guangxiense |
| Acinetobacter sp. NEB 394 | Acinetobacter sp. NEB 394 | Bradyrhizobium japonicum |
| Acinetobacter sp. SWBY1 | Acinetobacter sp. SWBY1 | Bradyrhizobium oligotrophicum |
| Acinetobacter sp. WCHA55 | Acinetobacter sp. WCHA55 | Bradyrhizobium sp. 1(2017) |
| Acinetobacter sp. WCHAc010034 | Acinetobacter sp. WCHAc010034 | Bradyrhizobium sp. 323S2 |
| Acinetobacter sp. WCHAc010052 | Acinetobacter sp. WCHAc010052 | Bradyrhizobium sp. BTAi1 |
| Acinetobacter tandoii | Acinetobacter tandoii | Bradyrhizobium sp. CCGE-LA001 |
| Acinetobacter towneri | Acinetobacter towneri | Bradyrhizobium sp. LCT2 |
| Acinetobacter ursingii | Acinetobacter ursingii | Bradyrhizobium sp. ORS 285 |
| Acinetobacter venetianus | Acinetobacter venetianus | Bradyrhizobium sp. |
| Acinetobacter wuhouensis | Acinetobacter wuhouensis | Bradyrhizobium symbiodeficiens |
| Acinetobacter | Acinetobacter | Bradyrhizobium vignae |
| Advenella mimigardefordensis | Advenella mimigardefordensis | Bradyrhizobium zhanjiangense |
| Aeromonas caviae | Aeromonas caviae | Burkholderiales |
| Aeromonas dhakensis | Aeromonas dhakensis | Calothrix parietina |
| Aeromonas encheleia | Aeromonas encheleia | Calothrix sp. NIES-3974 |
| Aeromonas hydrophila | Aeromonas hydrophila | Cellvibrio sp. KY-GH-1 |
| Aeromonas media | Aeromonas media | Cellvibrio sp. PSBB006 |
| Aeromonas rivipollensis | Aeromonas rivipollensis | Cellvibrio sp. PSBB023 |
| Aeromonas salmonicida | Aeromonas salmonicida | Chondrocystis sp. NIES-4102 |
| Aeromonas schubertii | Aeromonas schubertii | Chromatiaceae bacterium 2141T.STBD.0c.01a |
| Aeromonas sp. 1805 | Aeromonas sp. 1805 | Cupriavidus campinensis |
| Aeromonas sp. ASNIH1 | Aeromonas sp. ASNIH1 | Cupriavidus gilardii |
| Aeromonas sp. ASNIH5 | Aeromonas sp. ASNIH5 | Cupriavidus nantongensis |
| Aeromonas sp. CA23 | Aeromonas sp. CA23 | Cupriavidus neocaledonicus |
| Aeromonas sp. CU5 | Aeromonas sp. CU5 | Cupriavidus oxalaticus |
| Aeromonas veronii | Aeromonas veronii | Cupriavidus sp. USMAA2-4 |
| Aeromonas | Aeromonas | Cupriavidus taiwanensis |
| Afipia sp. GAS231 | Afipia sp. GAS231 | Cupriavidus |
| Agrobacterium fabrum | Agrobacterium fabrum | Cycloclasticus sp. P1 |
| Agrobacterium larrymoorei | Agrobacterium larrymoorei | Cycloclasticus sp. PY97N |
| Agrobacterium rhizogenes | Agrobacterium rhizogenes | Dechloromonas aromatica |
| Agrobacterium sp. H13-3 | Agrobacterium sp. H13-3 | Dechloromonas sp. HYN0024 |
| Agrobacterium sp. T29 | Agrobacterium sp. T29 | Desulfomicrobium baculatum |
| Agrobacterium tumefaciens complex | Agrobacterium tumefaciens complex | Desulfomicrobium orale |
| Agrobacterium tumefaciens | Agrobacterium tumefaciens | Desulfosarcina alkanivorans |
| Agrobacterium vitis | Agrobacterium vitis | Desulfovibrio gigas |
| Agrobacterium | Agrobacterium | Desulfurispirillum indicum |
| Alcanivorax borkumensis | Alcanivorax borkumensis | Diaphorobacter polyhydroxybutyrativorans |
| Allochromatium vinosum | Allochromatium vinosum | Ferrimonas balearica |
| Alphaproteobacteria | Alphaproteobacteria | gamma proteobacterium SS-5 |
| Aminobacter sp. MSH1 | Aminobacter sp. MSH1 | Geminocystis herdmanii |
| Ancylobacter pratisalsi | Ancylobacter pratisalsi | Geminocystis sp. NIES-3708 |
| Aquaspirillum sp. LM1 | Aquaspirillum sp. LM1 | Gemmata obscuriglobus |
| Asticcacaulis excentricus | Asticcacaulis excentricus | Halomicronema hongdechloris |
| Bdellovibrio exovorus | Bdellovibrio exovorus | Halomonas sp. Y2R2 |
| Bordetella bronchiseptica | Bordetella bronchiseptica | Hydrogenobacter sp. T-8 |
| Bordetella genomosp. 6 | Bordetella genomosp. 6 | Hydrogenobacter thermophilus |
| Bordetella parapertussis | Bordetella parapertussis | Hydrogenophaga pseudoflava |
| Bordetella | Bordetella | Hydrogenophaga sp. NH-16 |
| Bradyrhizobiaceae | Bradyrhizobiaceae | Hydrogenovibrio crunogenus |
| Bradyrhizobium erythrophlei | Bradyrhizobium erythrophlei | Hydrogenovibrio thermophilus |
| Bradyrhizobium lablabi | Bradyrhizobium lablabi | Hylemonella gracilis |
| Bradyrhizobium | Bradyrhizobium | Hyphomicrobium nitrativorans |
| Brenneria goodwinii | Brenneria goodwinii | Janthinobacterium sp. 1_2014MBL_MicDiv |
| Brenneria rubrifaciens | Brenneria rubrifaciens | Kiritimatiellaeota bacterium S-5007 |
| Brevundimonas diminuta | Brevundimonas diminuta | Leptolyngbya sp. O-77 |
| Brevundimonas sp. Bb-A | Brevundimonas sp. Bb-A | Lichenicola cladoniae |
| Brevundimonas sp. DS20 | Brevundimonas sp. DS20 | Luteithermobacter gelatinilyticus |
| Brevundimonas | Brevundimonas | Luteovulum sphaeroides |
| Burkholderia ambifaria | Burkholderia ambifaria | Magnetospira sp. QH-2 |
| Burkholderia anthina | Burkholderia anthina | Magnetospirillum magneticum |
| Burkholderia cenocepacia | Burkholderia cenocepacia | Magnetospirillum |
| Burkholderia cepacia | Burkholderia cepacia | Malaciobacter mytili |
| Burkholderia contaminans | Burkholderia contaminans | Marichromatium purpuratum |
| Burkholderia gladioli | Burkholderia gladioli | Marinobacter fonticola |
| Burkholderia glumae | Burkholderia glumae | Marinobacter salarius |
| Burkholderia lata | Burkholderia lata | Marinobacter |
| Burkholderia latens | Burkholderia latens | Methylibium petroleiphilum |
| Burkholderia multivorans | Burkholderia multivorans | Methylibium sp. Pch-M |
| Burkholderia oklahomensis | Burkholderia oklahomensis | Methylobacterium brachiatum |

| List names | Number of elements | Number of unique elements |
|---|---|---|
| Bacterial organism with AitP-like | 759 | 759 |
| Bacterial organism with FrmR-AitP-like | 475 | 475 |
| Bacterial organism with non-FrmR-AitP-like | 284 | 284 |
| **Overall number of unique elements** | | **759** |

| | | |
|---|---|---|
| Burkholderia plantarii | Burkholderia plantarii | Methylomicrobium alcaliphilum |
| Burkholderia pseudomallei | Burkholderia pseudomallei | Methylomicrobium buryatense |
| Burkholderia pseudomultivorans | Burkholderia pseudomultivorans | Methylomicrobium sp. wino1 |
| Burkholderia pyrrocinia | Burkholderia pyrrocinia | Methylomonas koyamae |
| Burkholderia seminalis | Burkholderia seminalis | Methylomonas methanica |
| Burkholderia sp. BDU6 | Burkholderia sp. BDU6 | Methylomonas rhizoryzae |
| Burkholderia sp. BDU8 | Burkholderia sp. BDU8 | Methylomonas sp. DH-1 |
| Burkholderia sp. DHOD12 | Burkholderia sp. DHOD12 | Methylomonas sp. LW13 |
| Burkholderia sp. IDO3 | Burkholderia sp. IDO3 | Methylophilus methylotrophus |
| Burkholderia sp. JP2-270 | Burkholderia sp. JP2-270 | Methyloversatilis sp. RAC08 |
| Burkholderia sp. KBS0801 | Burkholderia sp. KBS0801 | Methylovirgula ligni |
| Burkholderia sp. KK1 | Burkholderia sp. KK1 | Methylovulum psychrotolerans |
| Burkholderia sp. LA-2-3-30-S1-D2 | Burkholderia sp. LA-2-3-30-S1-D2 | Mitsuaria sp. 7 |
| Burkholderia sp. MSMB0856 | Burkholderia sp. MSMB0856 | Moritella yayanosii |
| Burkholderia sp. NRF60-BP8 | Burkholderia sp. NRF60-BP8 | Mucilaginibacter |
| Burkholderia sp. THE68 | Burkholderia sp. THE68 | Niabella ginsenosidivorans |
| Burkholderia stabilis | Burkholderia stabilis | Novosphingobium pentaromativorans |
| Burkholderia stagnalis | Burkholderia stagnalis | Oryzomicrobium terrae |
| Burkholderia territorii | Burkholderia territorii | Ottowia sp. oral taxon 894 |
| Burkholderia thailandensis | Burkholderia thailandensis | Paraburkholderia sprentiae |
| Burkholderia ubonensis | Burkholderia ubonensis | Phenylobacterium zucineum |
| Burkholderia | Burkholderia | Photobacterium gaetbulicola |
| Burkholderiaceae | Burkholderiaceae | Photobacterium profundum |
| Burkholderiales bacterium GJ-E10 | Burkholderiales bacterium GJ-E10 | Planctomycetes bacterium EC9 |
| Campylobacter hyointestinalis | Campylobacter hyointestinalis | Porphyrobacter sp. YT40 |
| Campylobacter iguaniorum | Campylobacter iguaniorum | Prolixibacteraceae bacterium WC007 |
| Campylobacter lanienae | Campylobacter lanienae | Pseudolabrys taiwanensis |
| Caulobacter mirabilis | Caulobacter mirabilis | Pseudomonadaceae bacterium SI-3 |
| Chelativorans | Chelativorans | Pseudomonas balearica |
| Chlorobium limicola | Chlorobium limicola | Pseudomonas fluorescens group |
| Collimonas pratensis | Collimonas pratensis | Pseudomonas fuscovaginae |
| Colwellia psychrerythraea | Colwellia psychrerythraea | Pseudomonas lalkuanensis |
| Colwellia sp. PAMC 20917 | Colwellia sp. PAMC 20917 | Pseudomonas marincola |
| Comamonadaceae | Comamonadaceae | Pseudomonas oryzae |
| Comamonas aquatica | Comamonas aquatica | Pseudomonas sp. CC6-YY-74 |
| Comamonas testosteroni | Comamonas testosteroni | Pseudomonas sp. LTJR-52 |
| Comamonas thiooxydans | Comamonas thiooxydans | Pseudomonas sp. R2A2 |
| Comamonas | Comamonas | Pseudomonas sp. THAF7b |
| Congregibacter litoralis | Congregibacter litoralis | Pseudomonas stutzeri |
| Croceicoccus naphthovorans | Croceicoccus naphthovorans | Pseudomonas xanthomarina |
| Cupriavidus basilensis | Cupriavidus basilensis | Pseudorhodoplanes sinuspersici |
| Cupriavidus metallidurans | Cupriavidus metallidurans | Psychromonas sp. CNPT3 |
| Cupriavidus necator | Cupriavidus necator | Qipengyuania flava |
| Cupriavidus pauculus | Cupriavidus pauculus | Ralstonia solanacearum |
| Cycloclasticus zancles | Cycloclasticus zancles | Raphidiopsis curvata |
| Delftia acidovorans | Delftia acidovorans | Rhodocista sp. MIMtkB3 |
| Delftia sp. Cs1-4 | Delftia sp. Cs1-4 | Rhodoferax sediminis |
| Delftia tsuruhatensis | Delftia tsuruhatensis | Rhodoplanes sp. Z2-YC6860 |
| Desulfococcus multivorans | Desulfococcus multivorans | Rhodopseudomonas palustris |
| Devosia ginsengisoli | Devosia ginsengisoli | Rhodospirillum centenum |
| Devosia sp. A16 | Devosia sp. A16 | Rhodospirillum rubrum |
| Diaphorobacter sp. HDW4A | Diaphorobacter sp. HDW4A | Rhodovulum sp. P5 |
| Diaphorobacter sp. JS3050 | Diaphorobacter sp. JS3050 | Rosistilla carotiformis |
| Ensifer alkalisoli | Ensifer alkalisoli | Rosistilla oblonga |
| Ensifer sojae | Ensifer sojae | Rubrivivax gelatinosus |
| Gammaproteobacteria | Gammaproteobacteria | Salinispira pacifica |
| Georhizobium profundi | Georhizobium profundi | Sedimenticola thiotaurini |
| Gibbsiella quercinecans | Gibbsiella quercinecans | Shewanella aestuarii |
| Herbaspirillum huttiense | Herbaspirillum huttiense | Shewanella algae |
| Herbaspirillum robiniae | Herbaspirillum robiniae | Shewanella amazonensis |
| Herbaspirillum rubrisubalbicans | Herbaspirillum rubrisubalbicans | Shewanella baltica |
| Herbaspirillum seropedicae | Herbaspirillum seropedicae | Shewanella benthica |
| Herbaspirillum | Herbaspirillum | Shewanella bicestrii |
| Herminiimonas arsenitoxidans | Herminiimonas arsenitoxidans | Shewanella chilikensis |
| Hydrogenophaga sp. BA0156 | Hydrogenophaga sp. BA0156 | Shewanella decolorationis |
| Hyphomicrobium denitrificans | Hyphomicrobium denitrificans | Shewanella donghaensis |
| Janthinobacterium agaricidamnosum | Janthinobacterium agaricidamnosum | Shewanella frigidimarina |
| Janthinobacterium lividum | Janthinobacterium lividum | Shewanella halifaxensis |
| Janthinobacterium sp. LM6 | Janthinobacterium sp. LM6 | Shewanella japonica |
| Janthinobacterium sp. Marseille | Janthinobacterium sp. Marseille | Shewanella khirikhana |
| Janthinobacterium sp. SNU WT3 | Janthinobacterium sp. SNU WT3 | Shewanella livingstonensis |
| Janthinobacterium svalbardensis | Janthinobacterium svalbardensis | Shewanella loihica |
| Ketogulonicigenium vulgare | Ketogulonicigenium vulgare | Shewanella marisflavi |
| Lautropia mirabilis | Lautropia mirabilis | Shewanella maritima |
| Lichenihabitans psoromatis | Lichenihabitans psoromatis | Shewanella oneidensis |
| Luteibacter pinisoli | Luteibacter pinisoli | Shewanella pealeana |
| Lysobacter soli | Lysobacter soli | Shewanella piezotolerans |
| Magnetospirillum gryphiswaldense | Magnetospirillum gryphiswaldense | Shewanella polaris |
| Marinobacter sp. BSs20148 | Marinobacter sp. BSs20148 | Shewanella psychrophila |
| Martelella endophytica | Martelella endophytica | Shewanella putrefaciens |
| Massilia flava | Massilia flava | Shewanella sediminis |
| Massilia putida | Massilia putida | Shewanella sp. ANA-3 |
| Mesorhizobium sp. NZP2077 | Mesorhizobium sp. NZP2077 | Shewanella sp. FDAARGOS_354 |
| Mesorhizobium | Mesorhizobium | Shewanella sp. MEBiC00475 |
| Methylomicrobium album | Methylomicrobium album | Shewanella sp. MR-4 |
| Microvirgula aerodenitrificans | Microvirgula aerodenitrificans | Shewanella sp. MR-7 |
| Moraxellaceae bacterium HYN0046 | Moraxellaceae bacterium HYN0046 | Shewanella sp. Pdp11 |
| Neoasaia chiangmaiensis | Neoasaia chiangmaiensis | Shewanella sp. Scap07 |
| Neorhizobium galegae | Neorhizobium galegae | Shewanella sp. W3-18-1 |
| Neorhizobium sp. NCHU2750 | Neorhizobium sp. NCHU2750 | Shewanella sp. WE21 |
| Nitrosomonas eutropha | Nitrosomonas eutropha | Shewanella sp. YLB-06 |
| Noviherbaspirillum sp. UKPF54 | Noviherbaspirillum sp. UKPF54 | Shewanella violacea |
| Novosphingobium aromaticivorans | Novosphingobium aromaticivorans | Shewanella woodyi |
| Novosphingobium resinovorum | Novosphingobium resinovorum | Shewanella |
| Novosphingobium sp. P6W | Novosphingobium sp. P6W | Sinorhizobium/Ensifer group |
| Novosphingobium sp. PP1Y | Novosphingobium sp. PP1Y | Skermanella pratensis |
| Novosphingobium tardaugens | Novosphingobium tardaugens | Sphaerospermopsis |
| Ochrobactrum | Ochrobactrum | Sphingobium amiense |
| Pandoraea pnomenusa | Pandoraea pnomenusa | Sphingobium fuliginis |
| Pandoraea sp. XY-2 | Pandoraea sp. XY-2 | Sphingobium sp. CAP-1 |
| Pannonibacter phragmitetus | Pannonibacter phragmitetus | Sphingobium sp. SCG-1 |
| Paraburkholderia acidiphila | Paraburkholderia acidiphila | Sphingomonas indica |
| Paraburkholderia acidisoli | Paraburkholderia acidisoli | Sphingomonas sp. C33 |
| Paraburkholderia acidophila | Paraburkholderia acidophila | Sphingomonas xanthus |
| Paraburkholderia aromaticivorans | Paraburkholderia aromaticivorans | Sphingopyxis fribergensis |
| Paraburkholderia atlantica | Paraburkholderia atlantica | Sphingopyxis sp. MG |
| Paraburkholderia caffeinilytica | Paraburkholderia caffeinilytica | Sphingopyxis sp. PAMC25046 |

| | | |
|---|---|---|
| Paraburkholderia fungorum | Paraburkholderia fungorum | Sphingopyxis sp. QXT-31 |
| Paraburkholderia phytofirmans | Paraburkholderia phytofirmans | Sphingosinicella sp. BN140058 |
| Paraburkholderia sp. 7MH5 | Paraburkholderia sp. 7MH5 | Stanieria cyanosphaera |
| Paraburkholderia xenovorans | Paraburkholderia xenovorans | Sulfuricaulis limicola |
| Paraburkholderia | Paraburkholderia | Synechococcus sp. PCC 6312 |
| Paracoccus denitrificans | Paracoccus denitrificans | Synechococcus sp. RSCCF101 |
| Paracoccus sp. AK26 | Paracoccus sp. AK26 | Synechocystis sp. PCC 6714 |
| Parvularcula bermudensis | Parvularcula bermudensis | Syntrophotalea acetylenivorans |
| Pelagibacterium halotolerans | Pelagibacterium halotolerans | Syntrophotalea carbinolica |
| Phreatobacter stygius | Phreatobacter stygius | Taylorella asinigenitalis |
| Phyllobacterium zundukense | Phyllobacterium zundukense | Taylorella equigenitalis |
| Pigmentiphaga sp. H8 | Pigmentiphaga sp. H8 | Terasakiella sp. SH-1 |
| Porphyrobacter sp. CACIAM 03H1 | Porphyrobacter sp. CACIAM 03H1 | Teredinibacter turnerae |
| Proteobacteria | Proteobacteria | Thauera sp. MZ1T |
| Pseudoarcobacter acticola | Pseudoarcobacter acticola | Thermocrinis albus |
| Pseudolysobacter antarcticus | Pseudolysobacter antarcticus | Thermocrinis ruber |
| pseudomallei group | pseudomallei group | Thiocystis violascens |
| Pseudomonas aeruginosa PAO1 | Pseudomonas aeruginosa PAO1 | Thioflavicoccus mobilis |
| Pseudomonas aeruginosa | Pseudomonas aeruginosa | Thiomicrorhabdus indica |
| Pseudomonas antarctica | Pseudomonas antarctica | Thiomicrospira sp. S5 |
| Pseudomonas arsenicoxydans | Pseudomonas arsenicoxydans | Thiospirochaeta perfilievii |
| Pseudomonas asplenii | Pseudomonas asplenii | Tistrella mobilis |
| Pseudomonas asturiensis | Pseudomonas asturiensis | Trichormus azollae |
| Pseudomonas cerasi | Pseudomonas cerasi | unclassified Azospirillum |
| Pseudomonas chlororaphis | Pseudomonas chlororaphis | unclassified Beijerinckiaceae |
| Pseudomonas corrugata | Pseudomonas corrugata | unclassified Bradyrhizobium |
| Pseudomonas denitrificans (nom. rej.) | Pseudomonas denitrificans (nom. rej.) | unclassified Caballeronia |
| Pseudomonas entomophila | Pseudomonas entomophila | unclassified Campylobacter |
| Pseudomonas extremaustralis | Pseudomonas extremaustralis | unclassified Halomonas |
| Pseudomonas fluorescens | Pseudomonas fluorescens | unclassified Methylobacterium |
| Pseudomonas frederiksbergensis | Pseudomonas frederiksbergensis | unclassified Planctomycetes |
| Pseudomonas knackmussii | Pseudomonas knackmussii | unclassified Shewanella |
| Pseudomonas koreensis | Pseudomonas koreensis | unclassified Synechococcus |
| Pseudomonas kribbensis | Pseudomonas kribbensis | unclassified Tardiphaga |
| Pseudomonas lini | Pseudomonas lini | unclassified Thermoleptolyngbya |
| Pseudomonas mandelii | Pseudomonas mandelii | unclassified Variovorax |
| Pseudomonas mediterranea | Pseudomonas mediterranea | unclassified Vibrio |
| Pseudomonas monteilii | Pseudomonas monteilii | Usitatibacter rugosus |
| Pseudomonas multiresinivorans | Pseudomonas multiresinivorans | Variovorax boronicumulans |
| Pseudomonas nitroreducens | Pseudomonas nitroreducens | Variovorax paradoxus |
| Pseudomonas palleroniana | Pseudomonas palleroniana | Variovorax sp. PAMC 28711 |
| Pseudomonas protegens | Pseudomonas protegens | Variovorax sp. PBL-H6 |
| Pseudomonas psychrotolerans | Pseudomonas psychrotolerans | Variovorax sp. PBS-H4 |
| Pseudomonas putida group | Pseudomonas putida group | Variovorax sp. RA8 |
| Pseudomonas putida | Pseudomonas putida | Verrucomicrobia bacterium S94 |
| Pseudomonas reinekei | Pseudomonas reinekei | Vibrio anguillarum |
| Pseudomonas sp. 09C 129 | Pseudomonas sp. 09C 129 | Vibrio astriarenae |
| Pseudomonas sp. 31-12 | Pseudomonas sp. 31-12 | Vibrio atlanticus |
| Pseudomonas sp. ABC1 | Pseudomonas sp. ABC1 | Vibrio campbellii |
| Pseudomonas sp. ACM7 | Pseudomonas sp. ACM7 | Vibrio coralliilyticus |
| Pseudomonas sp. ADAK18 | Pseudomonas sp. ADAK18 | Vibrio fluvialis |
| Pseudomonas sp. ATCC 13867 | Pseudomonas sp. ATCC 13867 | Vibrio furnissii |
| Pseudomonas sp. BIOMIG1BAC | Pseudomonas sp. BIOMIG1BAC | Vibrio harveyi group |
| Pseudomonas sp. BJP69 | Pseudomonas sp. BJP69 | Vibrio harveyi |
| Pseudomonas sp. CMR12a | Pseudomonas sp. CMR12a | Vibrio hyugaensis |
| Pseudomonas sp. CMR5c | Pseudomonas sp. CMR5c | Vibrio mediterranei |
| Pseudomonas sp. DTU12.3 | Pseudomonas sp. DTU12.3 | Vibrio natriegens |
| Pseudomonas sp. FGI182 | Pseudomonas sp. FGI182 | Vibrio owensii |
| Pseudomonas sp. IB20 | Pseudomonas sp. IB20 | Vibrio panuliri |
| Pseudomonas sp. J380 | Pseudomonas sp. J380 | Vibrio qinghaiensis |
| Pseudomonas sp. JY-Q | Pseudomonas sp. JY-Q | Vibrio rotiferianus |
| Pseudomonas sp. K2W31S-8 | Pseudomonas sp. K2W31S-8 | Vibrio sp. dhg |
| Pseudomonas sp. Leaf58 | Pseudomonas sp. Leaf58 | Vibrio sp. EJY3 |
| Pseudomonas sp. MRSN12121 | Pseudomonas sp. MRSN12121 | Vibrio sp. THAF190c |
| Pseudomonas sp. NIBRBAC000502773 | Pseudomonas sp. NIBRBAC000502773 | Vibrio sp. ZWAL4003 |
| Pseudomonas sp. NP-1 | Pseudomonas sp. NP-1 | Vibrio splendidus |
| Pseudomonas sp. R11-23-07 | Pseudomonas sp. R11-23-07 | Vibrio tapetis |
| Pseudomonas sp. R76 | Pseudomonas sp. R76 | Vibrio tritonius |
| Pseudomonas sp. S150 | Pseudomonas sp. S150 | Vibrio vulnificus |
| Pseudomonas sp. SCB32 | Pseudomonas sp. SCB32 | Vibrio zhugei |
| Pseudomonas sp. SK | Pseudomonas sp. SK | Vitreoscilla filiformis |
| Pseudomonas sp. StFLB209 | Pseudomonas sp. StFLB209 | Xanthomonas albilineans |
| Pseudomonas sp. SWI6 | Pseudomonas sp. SWI6 | Xanthomonas sacchari |
| Pseudomonas sp. TCU-HL1 | Pseudomonas sp. TCU-HL1 | Xylophilus rhododendri |
| Pseudomonas sp. TUM18999 | Pseudomonas sp. TUM18999 | |
| Pseudomonas sp. UW4 | Pseudomonas sp. UW4 | |
| Pseudomonas syringae group | Pseudomonas syringae group | |
| Pseudomonas syringae | Pseudomonas syringae | |
| Pseudomonas umsongensis | Pseudomonas umsongensis | |
| Pseudomonas vancouverensis | Pseudomonas vancouverensis | |
| Pseudomonas versuta | Pseudomonas versuta | |
| Pseudomonas viridiflava | Pseudomonas viridiflava | |
| Pseudomonas xinjiangensis | Pseudomonas xinjiangensis | |
| Pseudomonas yamanorum | Pseudomonas yamanorum | |
| Pseudomonas | Pseudomonas | |
| Pseudoxanthomonas spadix | Pseudoxanthomonas spadix | |
| Pusillimonas sp. T7-7 | Pusillimonas sp. T7-7 | |
| Qipengyuania sediminis | Qipengyuania sediminis | |
| Ralstonia pickettii | Ralstonia pickettii | |
| Ralstonia | Ralstonia | |
| Rhizobacter gummiphilus | Rhizobacter gummiphilus | |
| Rhizobiaceae | Rhizobiaceae | |
| Rhizobiales | Rhizobiales | |
| Rhizobium acidisoli | Rhizobium acidisoli | |
| Rhizobium daejeonense | Rhizobium daejeonense | |
| Rhizobium etli | Rhizobium etli | |
| Rhizobium favelukesii | Rhizobium favelukesii | |
| Rhizobium gallicum | Rhizobium gallicum | |
| Rhizobium jaguaris | Rhizobium jaguaris | |
| Rhizobium leguminosarum | Rhizobium leguminosarum | |
| Rhizobium phaseoli | Rhizobium phaseoli | |
| Rhizobium pusense | Rhizobium pusense | |
| Rhizobium rhizoryzae | Rhizobium rhizoryzae | |
| Rhizobium sp. 11515TR | Rhizobium sp. 11515TR | |
| Rhizobium sp. ACO-34A | Rhizobium sp. ACO-34A | |
| Rhizobium sp. CIAT894 | Rhizobium sp. CIAT894 | |
| Rhizobium sp. NIBRBAC000502774 | Rhizobium sp. NIBRBAC000502774 | |

| | |
|---|---|
| Rhizobium sp. NXC14 | Rhizobium sp. NXC14 |
| Rhizobium sp. NXC24 | Rhizobium sp. NXC24 |
| Rhizobium/Agrobacterium group | Rhizobium/Agrobacterium group |
| Rhizobium | Rhizobium |
| Rhodanobacter | Rhodanobacter |
| Rhodobacter capsulatus | Rhodobacter capsulatus |
| Rhodobacter sp. CZR27 | Rhodobacter sp. CZR27 |
| Rhodovulum sp. MB263 | Rhodovulum sp. MB263 |
| Rhodovulum sulfidophilum | Rhodovulum sulfidophilum |
| Roseateles depolymerans | Roseateles depolymerans |
| Roseomonas sp. FDAARGOS_362 | Roseomonas sp. FDAARGOS_362 |
| Roseomonas | Roseomonas |
| Salinimonas sediminis | Salinimonas sediminis |
| Serratia marcescens | Serratia marcescens |
| Serratia odorifera | Serratia odorifera |
| Serratia sp. FS14 | Serratia sp. FS14 |
| Serratia surfactantfaciens | Serratia surfactantfaciens |
| Serratia | Serratia |
| Shinella sp. HZN7 | Shinella sp. HZN7 |
| Simiduia agarivorans | Simiduia agarivorans |
| Sinorhizobium fredii | Sinorhizobium fredii |
| Sinorhizobium meliloti | Sinorhizobium meliloti |
| Sinorhizobium | Sinorhizobium |
| Sorangium cellulosum | Sorangium cellulosum |
| Sphingobium baderi | Sphingobium baderi |
| Sphingobium chlorophenolicum | Sphingobium chlorophenolicum |
| Sphingobium cloacae | Sphingobium cloacae |
| Sphingobium herbicidovorans | Sphingobium herbicidovorans |
| Sphingobium hydrophobicum | Sphingobium hydrophobicum |
| Sphingobium japonicum | Sphingobium japonicum |
| Sphingobium sp. EP60837 | Sphingobium sp. EP60837 |
| Sphingobium sp. MI1205 | Sphingobium sp. MI1205 |
| Sphingobium sp. RAC03 | Sphingobium sp. RAC03 |
| Sphingobium sp. RSMS | Sphingobium sp. RSMS |
| Sphingobium yanoikuyae | Sphingobium yanoikuyae |
| Sphingobium | Sphingobium |
| Sphingomonadaceae | Sphingomonadaceae |
| Sphingomonas hengshuiensis | Sphingomonas hengshuiensis |
| Sphingomonas sp. C8-2 | Sphingomonas sp. C8-2 |
| Sphingomonas sp. IC081 | Sphingomonas sp. IC081 |
| Sphingomonas sp. KC8 | Sphingomonas sp. KC8 |
| Sphingomonas sp. LK11 | Sphingomonas sp. LK11 |
| Sphingomonas wittichii | Sphingomonas wittichii |
| Sphingomonas | Sphingomonas |
| Sphingopyxis alaskensis | Sphingopyxis alaskensis |
| Sphingopyxis granuli | Sphingopyxis granuli |
| Sphingopyxis lindanitolerans | Sphingopyxis lindanitolerans |
| Sphingopyxis macrogoltabida | Sphingopyxis macrogoltabida |
| Sphingopyxis sp. 113P3 | Sphingopyxis sp. 113P3 |
| Sphingopyxis sp. FD7 | Sphingopyxis sp. FD7 |
| Sphingopyxis | Sphingopyxis |
| Sphingorhabdus sp. SMR4y | Sphingorhabdus sp. SMR4y |
| Sphingorhabdus sp. YGSMI21 | Sphingorhabdus sp. YGSMI21 |
| Sphingosinicella microcystinivorans | Sphingosinicella microcystinivorans |
| Stappia indica | Stappia indica |
| Starkeya sp. ORNL1 | Starkeya sp. ORNL1 |
| Stenotrophomonas acidaminiphila | Stenotrophomonas acidaminiphila |
| Stenotrophomonas indicatrix | Stenotrophomonas indicatrix |
| Stenotrophomonas maltophilia group | Stenotrophomonas maltophilia group |
| Stenotrophomonas maltophilia | Stenotrophomonas maltophilia |
| Stenotrophomonas sp. 364 | Stenotrophomonas sp. 364 |
| Stenotrophomonas sp. ASS1 | Stenotrophomonas sp. ASS1 |
| Stenotrophomonas sp. ESTM1D_MKCIP4_1 | Stenotrophomonas sp. ESTM1D_MKCIP4_1 |
| Stenotrophomonas sp. G4 | Stenotrophomonas sp. G4 |
| Stenotrophomonas sp. LM091 | Stenotrophomonas sp. LM091 |
| Stenotrophomonas sp. MYb57 | Stenotrophomonas sp. MYb57 |
| Stenotrophomonas sp. NA06056 | Stenotrophomonas sp. NA06056 |
| Stenotrophomonas sp. SAU14A_NAIMI4_8 | Stenotrophomonas sp. SAU14A_NAIMI4_8 |
| Stenotrophomonas sp. WZN-1 | Stenotrophomonas sp. WZN-1 |
| Stenotrophomonas sp. YAU14D1_LEIMI4_1 | Stenotrophomonas sp. YAU14D1_LEIMI4_1 |
| Stenotrophomonas sp. ZAC14D2_NAIMI4_6 | Stenotrophomonas sp. ZAC14D2_NAIMI4_6 |
| Stenotrophomonas | Stenotrophomonas |
| Sulfuricurvum kujiense | Sulfuricurvum kujiense |
| Sulfuriferula nivalis | Sulfuriferula nivalis |
| Sulfurospirillum cavolei | Sulfurospirillum cavolei |
| Sulfurospirillum halorespirans | Sulfurospirillum halorespirans |
| Sulfurospirillum multivorans | Sulfurospirillum multivorans |
| Tateyamaria omphalii | Tateyamaria omphalii |
| Thalassococcus sp. S3 | Thalassococcus sp. S3 |
| Thalassolituus oleivorans | Thalassolituus oleivorans |
| Thauera aromatica | Thauera aromatica |
| Thauera sp. K11 | Thauera sp. K11 |
| Thauera | Thauera |
| Thiomonas sp. X19 | Thiomonas sp. X19 |
| Thiomonas | Thiomonas |
| Tolumonas auensis | Tolumonas auensis |
| unclassified Agrobacterium | unclassified Agrobacterium |
| unclassified Burkholderia | unclassified Burkholderia |
| unclassified Marinobacter | unclassified Marinobacter |
| unclassified Pseudomonas | unclassified Pseudomonas |
| unclassified Rhizobium | unclassified Rhizobium |
| unclassified Roseomonas | unclassified Roseomonas |
| unclassified Sphingobium | unclassified Sphingobium |
| unclassified Thiomonas | unclassified Thiomonas |
| Undibacterium sp. KW1 | Undibacterium sp. KW1 |
| Variovorax sp. HW608 | Variovorax sp. HW608 |
| Variovorax sp. PBL-E5 | Variovorax sp. PBL-E5 |
| Variovorax sp. SRS16 | Variovorax sp. SRS16 |
| Vibrio parahaemolyticus | Vibrio parahaemolyticus |
| Vibrio | Vibrio |
| Xanthomonas arboricola | Xanthomonas arboricola |
| Xanthomonas axonopodis | Xanthomonas axonopodis |
| Xanthomonas campestris | Xanthomonas campestris |
| Xanthomonas cassavae | Xanthomonas cassavae |
| Xanthomonas citri | Xanthomonas citri |
| Xanthomonas cucurbitae | Xanthomonas cucurbitae |
| Xanthomonas euvesicatoria | Xanthomonas euvesicatoria |

| | |
|---|---|
| Xanthomonas hortorum | Xanthomonas hortorum |
| Xanthomonas oryzae | Xanthomonas oryzae |
| Xanthomonas phaseoli | Xanthomonas phaseoli |
| Xanthomonas vasicola | Xanthomonas vasicola |
| Xanthomonas vesicatoria | Xanthomonas vesicatoria |
| Xanthomonas | Xanthomonas |
| Zhongshania aliphaticivorans | Zhongshania aliphaticivorans |
| Zymomonas mobilis | Zymomonas mobilis |
| Acinetobacter oleivorans CIP 110421 | Acinetobacter oleivorans CIP 110421 |
| Aeromonas sp. 2692-1 | Aeromonas sp. 2692-1 |
| Azoarcus sp. KH32C | Azoarcus sp. KH32C |
| Azospirillum humicireducens | Azospirillum humicireducens |
| Bosea sp. AS-1 | Bosea sp. AS-1 |
| Bosea sp. F3-2 | Bosea sp. F3-2 |
| Burkholderia diffusa | Burkholderia diffusa |
| Burkholderia vietnamiensis | Burkholderia vietnamiensis |
| Desulfatibacillum aliphaticivorans | Desulfatibacillum aliphaticivorans |
| Halomonas piezotolerans | Halomonas piezotolerans |
| Halomonas sp. hl-4 | Halomonas sp. hl-4 |
| Halomonas subglaciescola | Halomonas subglaciescola |
| Halomonas titanicae | Halomonas titanicae |
| Hypericibacter adhaerens | Hypericibacter adhaerens |
| Hypericibacter terrae | Hypericibacter terrae |
| Hyphomicrobium nitrativorans NL23 | Hyphomicrobium nitrativorans NL23 |
| Indioceanicola profundi | Indioceanicola profundi |
| Rhodobacter sphaeroides ATCC 17029 | Rhodobacter sphaeroides ATCC 17029 |
| Malaciobacter mytili LMG 24559 | Malaciobacter mytili LMG 24559 |
| Marinobacter sp. Arc7-DN-1 | Marinobacter sp. Arc7-DN-1 |
| Marinobacter hydrocarbonoclasticus VT8 | Marinobacter hydrocarbonoclasticus VT8 |
| Mesorhizobium sp. DCY119 | Mesorhizobium sp. DCY119 |
| Methylophilus methylotrophus DSM 46235 = AT | Methylophilus methylotrophus DSM 46235 = ATCC 53528 |
| Paracoccus liaowanqingii | Paracoccus liaowanqingii |
| Pseudomonas fuscovaginae CB98818 | Pseudomonas fuscovaginae CB98818 |
| Pseudomonas sp. R2-7-07 | Pseudomonas sp. R2-7-07 |
| Pseudomonas sp. R4-34-07 | Pseudomonas sp. R4-34-07 |
| Pseudomonas sp. SWI44 | Pseudomonas sp. SWI44 |
| Rhizobium sp. IE4771 | Rhizobium sp. IE4771 |
| Ensifer adhaerens | Ensifer adhaerens |
| Sphingobium fuliginis ATCC 27551 | Sphingobium fuliginis ATCC 27551 |
| Sphingobium sp. PAMC28499 | Sphingobium sp. PAMC28499 |
| Stenotrophomonas sp. SAU14A_NAIMI4_5 | Stenotrophomonas sp. SAU14A_NAIMI4_5 |
| Stenotrophomonas sp. YAU14A_MKIMI4_1 | Stenotrophomonas sp. YAU14A_MKIMI4_1 |
| Stenotrophomonas sp. ZAC14A_NAIMI4_1 | Stenotrophomonas sp. ZAC14A_NAIMI4_1 |
| Stenotrophomonas sp. ZAC14D2_NAIMI4_7 | Stenotrophomonas sp. ZAC14D2_NAIMI4_7 |
| Pelobacter carbinolicus DSM 2380 | Pelobacter carbinolicus DSM 2380 |
| Azospirillum sp. TSA2s | Azospirillum sp. TSA2s |
| Caballeronia sp. SBC1 | Caballeronia sp. SBC1 |
| Tardiphaga sp. vice154 | Tardiphaga sp. vice154 |
| Rhizobium pseudoryzae | Rhizobium pseudoryzae |
| Diaphorobacter sp. HDW4B | Diaphorobacter sp. HDW4B |
| Bradyrhizobium ottawaense | Bradyrhizobium ottawaense |
| Acaryochloris marina | |
| Acidihalobacter prosperus | |
| Acidovorax ebreus | |
| Acidovorax sp. KKS102 | |
| Acinetobacter calcoaceticus/baumannii complex | |
| Acinetobacter calcoaceticus | |
| Acinetobacter guillouiae | |
| Acinetobacter oleivorans | |
| Advenella kashmirensis | |
| Algicoccus marinus | |
| Altererythrobacter atlanticus | |
| Altererythrobacter sp. B11 | |
| Anabaena cylindrica | |
| Anaeromyxobacter dehalogenans | |
| Anaeromyxobacter sp. Fw109-5 | |
| Anaeromyxobacter sp. K | |
| Arcobacter cibarius | |
| Arcobacter nitrofigilis | |
| Azoarcus sp. CIB | |
| Azospirillum | |
| Azotobacter chroococcum | |
| Azotobacter salinestris | |
| Beggiatoa leptomitoformis | |
| Betaproteobacteria bacterium GR16-43 | |
| Blastochloris tepida | |
| Blastomonas fulva | |
| Blastomonas sp. RAC04 | |
| Bradyrhizobium amphicarpaeae | |
| Bradyrhizobium betae | |
| Bradyrhizobium canariense | |
| Bradyrhizobium cosmicum | |
| Bradyrhizobium diazoefficiens | |
| Bradyrhizobium guangxiense | |
| Bradyrhizobium japonicum | |
| Bradyrhizobium oligotrophicum | |
| Bradyrhizobium sp. 1(2017) | |
| Bradyrhizobium sp. 323S2 | |
| Bradyrhizobium sp. BTAi1 | |
| Bradyrhizobium sp. CCGE-LA001 | |
| Bradyrhizobium sp. LCT2 | |
| Bradyrhizobium sp. ORS 285 | |
| Bradyrhizobium sp. | |
| Bradyrhizobium symbiodeficiens | |
| Bradyrhizobium vignae | |
| Bradyrhizobium zhanjiangense | |
| Burkholderiales | |
| Calothrix parietina | |
| Calothrix sp. NIES-3974 | |
| Cellvibrio sp. KY-GH-1 | |
| Cellvibrio sp. PSBB006 | |
| Cellvibrio sp. PSBB023 | |
| Chondrocystis sp. NIES-4102 | |
| Chromatiaceae bacterium 2141T.STBD.0c.01a | |
| Cupriavidus campinensis | |
| Cupriavidus gilardii | |
| Cupriavidus nantongensis | |

Cupriavidus neocaledonicus
Cupriavidus oxalaticus
Cupriavidus sp. USMAA2-4
Cupriavidus taiwanensis
Cupriavidus
Cycloclasticus sp. P1
Cycloclasticus sp. PY97N
Dechloromonas aromatica
Dechloromonas sp. HYN0024
Desulfomicrobium baculatum
Desulfomicrobium orale
Desulfosarcina alkanivorans
Desulfovibrio gigas
Desulfurispirillum indicum
Diaphorobacter polyhydroxybutyrativorans
Ferrimonas balearica
gamma proteobacterium SS-5
Geminocystis herdmanii
Geminocystis sp. NIES-3708
Gemmata obscuriglobus
Halomicronema hongdechloris
Halomonas sp. Y2R2
Hydrogenobacter sp. T-8
Hydrogenobacter thermophilus
Hydrogenophaga pseudoflava
Hydrogenophaga sp. NH-16
Hydrogenovibrio crunogenus
Hydrogenovibrio thermophilus
Hylemonella gracilis
Hyphomicrobium nitrativorans
Janthinobacterium sp. 1_2014MBL_MicDiv
Kiritimatiellaeota bacterium S-5007
Leptolyngbya sp. O-77
Lichenicola cladoniae
Luteithermobacter gelatinilyticus
Luteovulum sphaeroides
Magnetospira sp. QH-2
Magnetospirillum magneticum
Magnetospirillum
Malaciobacter mytili
Marichromatium purpuratum
Marinobacter fonticola
Marinobacter salarius
Marinobacter
Methylibium petroleiphilum
Methylibium sp. Pch-M
Methylobacterium brachiatum
Methylomicrobium alcaliphilum
Methylomicrobium buryatense
Methylomicrobium sp. wino1
Methylomonas koyamae
Methylomonas methanica
Methylomonas rhizoryzae
Methylomonas sp. DH-1
Methylomonas sp. LW13
Methylophilus methylotrophus
Methyloversatilis sp. RAC08
Methylovirgula ligni
Methylovulum psychrotolerans
Mitsuaria sp. 7
Moritella yayanosii
Mucilaginibacter
Niabella ginsenosidivorans
Novosphingobium pentaromativorans
Oryzomicrobium terrae
Ottowia sp. oral taxon 894
Paraburkholderia sprentiae
Phenylobacterium zucineum
Photobacterium gaetbulicola
Photobacterium profundum
Planctomycetes bacterium EC9
Porphyrobacter sp. YT40
Prolixibacteraceae bacterium WC007
Pseudolabrys taiwanensis
Pseudomonadaceae bacterium SI-3
Pseudomonas balearica
Pseudomonas fluorescens group
Pseudomonas fuscovaginae
Pseudomonas lalkuanensis
Pseudomonas marincola
Pseudomonas oryzae
Pseudomonas sp. CC6-YY-74
Pseudomonas sp. LTJR-52
Pseudomonas sp. R2A2
Pseudomonas sp. THAF7b
Pseudomonas stutzeri
Pseudomonas xanthomarina
Pseudorhodoplanes sinuspersici
Psychromonas sp. CNPT3
Qipengyuania flava
Ralstonia solanacearum
Raphidiopsis curvata
Rhodocista sp. MIMtkB3
Rhodoferax sediminis
Rhodoplanes sp. Z2-YC6860
Rhodopseudomonas palustris
Rhodospirillum centenum
Rhodospirillum rubrum
Rhodovulum sp. P5
Rosistilla carotiformis
Rosistilla oblonga
Rubrivivax gelatinosus
Salinispira pacifica
Sedimenticola thiotaurini
Shewanella aestuarii
Shewanella algae
Shewanella amazonensis

| |
|---|
| Shewanella baltica |
| Shewanella benthica |
| Shewanella bicestrii |
| Shewanella chilikensis |
| Shewanella decolorationis |
| Shewanella donghaensis |
| Shewanella frigidimarina |
| Shewanella halifaxensis |
| Shewanella japonica |
| Shewanella khirikhana |
| Shewanella livingstonensis |
| Shewanella loihica |
| Shewanella marisflavi |
| Shewanella maritima |
| Shewanella oneidensis |
| Shewanella pealeana |
| Shewanella piezotolerans |
| Shewanella polaris |
| Shewanella psychrophila |
| Shewanella putrefaciens |
| Shewanella sediminis |
| Shewanella sp. ANA-3 |
| Shewanella sp. FDAARGOS_354 |
| Shewanella sp. MEBiC00475 |
| Shewanella sp. MR-4 |
| Shewanella sp. MR-7 |
| Shewanella sp. Pdp11 |
| Shewanella sp. Scap07 |
| Shewanella sp. W3-18-1 |
| Shewanella sp. WE21 |
| Shewanella sp. YLB-06 |
| Shewanella violacea |
| Shewanella woodyi |
| Shewanella |
| Sinorhizobium/Ensifer group |
| Skermanella pratensis |
| Sphaerospermopsis |
| Sphingobium amiense |
| Sphingobium fuliginis |
| Sphingobium sp. CAP-1 |
| Sphingobium sp. SCG-1 |
| Sphingomonas indica |
| Sphingomonas sp. C33 |
| Sphingomonas xanthus |
| Sphingopyxis fribergensis |
| Sphingopyxis sp. MG |
| Sphingopyxis sp. PAMC25046 |
| Sphingopyxis sp. QXT-31 |
| Sphingosinicella sp. BN140058 |
| Stanieria cyanosphaera |
| Sulfuricaulis limicola |
| Synechococcus sp. PCC 6312 |
| Synechococcus sp. RSCCF101 |
| Synechocystis sp. PCC 6714 |
| Syntrophotalea acetylenivorans |
| Syntrophotalea carbinolica |
| Taylorella asinigenitalis |
| Taylorella equigenitalis |
| Terasakiella sp. SH-1 |
| Teredinibacter turnerae |
| Thauera sp. MZ1T |
| Thermocrinis albus |
| Thermocrinis ruber |
| Thiocystis violascens |
| Thioflavicoccus mobilis |
| Thiomicrorhabdus indica |
| Thiomicrospira sp. S5 |
| Thiospirochaeta perfilievii |
| Tistrella mobilis |
| Trichormus azollae |
| unclassified Azospirillum |
| unclassified Beijerinckiaceae |
| unclassified Bradyrhizobium |
| unclassified Caballeronia |
| unclassified Campylobacter |
| unclassified Halomonas |
| unclassified Methylobacterium |
| unclassified Planctomycetes |
| unclassified Shewanella |
| unclassified Synechococcus |
| unclassified Tardiphaga |
| unclassified Thermoleptolyngbya |
| unclassified Variovorax |
| unclassified Vibrio |
| Usitatibacter rugosus |
| Variovorax boronicumulans |
| Variovorax paradoxus |
| Variovorax sp. PAMC 28711 |
| Variovorax sp. PBL-H6 |
| Variovorax sp. PBS-H4 |
| Variovorax sp. RA8 |
| Verrucomicrobia bacterium S94 |
| Vibrio anguillarum |
| Vibrio astriarenae |
| Vibrio atlanticus |
| Vibrio campbellii |
| Vibrio coralliilyticus |
| Vibrio fluvialis |
| Vibrio furnissii |
| Vibrio harveyi group |
| Vibrio harveyi |
| Vibrio hyugaensis |
| Vibrio mediterranei |
| Vibrio natriegens |
| Vibrio owensii |
| Vibrio panuliri |
| Vibrio qinghaiensis |

| |
|---|
| Vibrio rotiferianus |
| Vibrio sp. dhg |
| Vibrio sp. EJY3 |
| Vibrio sp. THAF190c |
| Vibrio sp. ZWAL4003 |
| Vibrio splendidus |
| Vibrio tapetis |
| Vibrio tritonius |
| Vibrio vulnificus |
| Vibrio zhugei |
| Vitreoscilla filiformis |
| Xanthomonas albilineans |
| Xanthomonas sacchari |
| Xylophilus rhododendri |
